# Influencer in flies: Socially interactive individuals shape group-level characteristics

**DOI:** 10.1101/2025.11.28.691157

**Authors:** Kaiya Hamamichi, Yuma Takahashi

## Abstract

Group living is a widespread adaptive strategy observed across the animal kingdom. Social interactions between individuals within a group often promote behavioral conformity, forming group-level characteristics that correspond to the group’s phenotypic and genotypic composition. However, for large-scale groups observed in non-social invertebrates, a theoretical framework establishing how genotypic composition influences group-level characteristics and their intergroup variation has not been verified. Here, we elucidated the mechanism governing activity conformity in multiple *Drosophila* strains. Furthermore, by combining various strains, we revealed the mechanism by which group-level characteristics are formed as a function of genotype composition. In single-strain groups, image analysis and causal analysis based on information theory revealed that all strains conformed their walking behavior to others’ behavior. In contrast, mixed-strain groups exhibited emergent group-level characteristics, with some individuals from certain strains displaying non-conforming behavior. Interestingly, these individuals exhibited high sociality and exerted strong social influence on others, suggesting that specific individuals (genotypes) acting as “influencers” within the group plays a crucial role in shaping group-level characteristics. In the wild, each group may develop phenotypic variation influenced by the presence of these individuals.

## Introduction

Group living is a widespread strategy across the animal kingdom, from insects to primates (1, 2). Sociality in a group brings clear benefits such as enhanced predator avoidance (3), improved foraging efficiency (4), and opportunities for learning from others (5). Learning from the behavior of other individuals and changing one’s own behavior to conform to theirs is called social learning (specifically, conformist social learning), and numerous cases have been reported (6, 7). For instance, where and how *Drosophila* flies forage (8), choose mates (9), or lay eggs are often influenced by the behavior of others through the mechanism of social learning (10), with direct consequences for fitness. These observations underscore the idea that the expression of individual phenotypes is not solely intrinsic, but arises within a social context. In other words, final behavioral phenotypes are determined through both one’s own genotype (Direct Genetic Effects: DGEs) and the genotypes of other individuals they interact with (Indirect Genetic Effects: IGEs) (11, 12). Therefore, to clarify how DGEs and IGEs shape behavioral variation among individuals in the wild, it is essential to elucidate the processes through which group-level characteristics (or performance) and their variation arise (13).

Conformity refers to an adjustment of one’s behavior in response to the behavior of others (14). This behavior represents a fundamental component of social learning and a key mechanism underlying IGEs (15). The modification of one’s own behavior in response to others’ behavior reduces behavioral variation within a group, ultimately causing all members to converge on a uniform phenotype (16). Such phenotypic convergence within groups occurs not only in behavior but also in resource use, metabolism, and physiological traits via behavioral changes, resulting in group members’ phenotypic convergence either immediately or incrementally via behavioral changes (17). These phenomena support the plausible theory that genotypic composition influences individual phenotypes. However, in non-social species forming non-moving groups (18), such as many invertebrates, if the group size is large enough that probabilistic phenotypic bias can be neglected, intergroup variation among groups with different genotypic compositions may disappear, leading to convergence toward similar phenotypes. In other words, it remains unclear how the genotypic composition of a group determines group-level characteristics through conformity in invertebrates (19).

Clarifying the relationship between genotypic composition within a group and the resulting convergent phenotypes leads to a better understanding of how DGEs and IGEs shape group-level characteristics, and opens the door to elucidating the contribution of genetic diversity to behavioral variation in wild populations (20–22).

The converged phenotypes resulting from conformity in a group were expected to align with the mean of members’ phenotypes (synthetic group-level traits) (23). On the other hand, behavioral variation among individuals and the degree of social interactions within a group shape intrinsic group-level traits (i.e., group-level characteristics) (24–26). For instance, an individual that interacts with many others has numerous opportunities to transfer its own phenotype to others through conformity, thereby largely contributing to the formation of group-level characteristics. In addition, if such individuals possess a tendency to take non-conforming other group members, the group-level characteristics will be more strongly influenced by such individuals (27).

That is, depending on its behavioral property, a part of individuals may function as an “influencer” within the group. Importantly, the presence of specific individuals (genotypes) could play an important role in shaping group-level characteristics, potentially contributing to the formation of intergroup phenotypic variation depending on the composition of group members (28).

*Drosophila* flies exhibit strong sociality (29, 30), and individual behavior is shaped by both social factors and IGEs (31, 32). Together with the availability of well-established behavioral assays (33, 34), *Drosophila* they provide an excellent system to study the relationship between genotypic composition and the resulting phenotype convergence in groups. In the present study, we assembled groups exhaustively by pairing multiple genotypes in a binary mixture design and examined both changes in individual-level locomotor behavior within the groups and shifts in the composition of group-level behavioral phenotypes in *Drosophila*, in order to identify social and behavioral mechanisms underlying how group-level characteristics are shaped.

## Results

To determine whether flies conform their activity to other individuals and identify the behavioral rules governing this conformity, we placed 24 individuals from each strain into a single arena, defined here as a “group”, and observed their behavior (Fig 1). The mean locomotive speed differed among the 20 strains (*df* = 110, χ^2^ = 562, *P* < 0.001), but remained constant over time (Fig. 2a). The variance components analysis showed that strain accounted for 26.3% of the variation in locomotive speed. In contrast, the effect of the group on locomotive speed variation was 37.1%, indicating that intergroup variation was larger than inter-strain variation. The within-group variation in locomotive speed observed for each strain was standardized by taking into account the variation in locomotive speed within virtual groups created by randomizing individuals across different arenas (groups) for each strain, and the standardized values of actual groups were negative for all strains, with the majority falling below a *z*-score of −1.96, corresponding to the lower 2.5% of the distribution (Fig. 2b). These results indicated that the locomotive speeds of individuals within actual groups converged toward similar values, suggesting phenotypic convergence. To clarify the causality between the total movement of other individuals (visual cue) and the focal individual’s locomotive speed, we quantified transfer entropy (TE). Results showed that the number of individuals showing clarity TE values (the degree of information transfer) was highest at the smallest lag time, and standardized TE values were consistently highest for the 180° field of view (Fig. 2c), indicating that an individual’s walking and stopping are influenced by fluctuations in visual cues in the forward visual field. In the subsequent analyses, we focused on the visual cues within a 180° field of view and a lag time of 0.5 seconds. The visual cue increased immediately before the onset of walking compared with stationary periods not associated with walking initiation for all strains (*df* = 3430, *t* = 52.3, *P* < 0.001), indicating that flies conformed their locomotive speed in response to the locomotive behavior of other individuals. Interestingly, visual cues were also higher just before stopping than during walking not associated with stopping (*df* = 3435, *t* = 33.9, *P* < 0.001; Fig. 2d).

**Figure 1.**
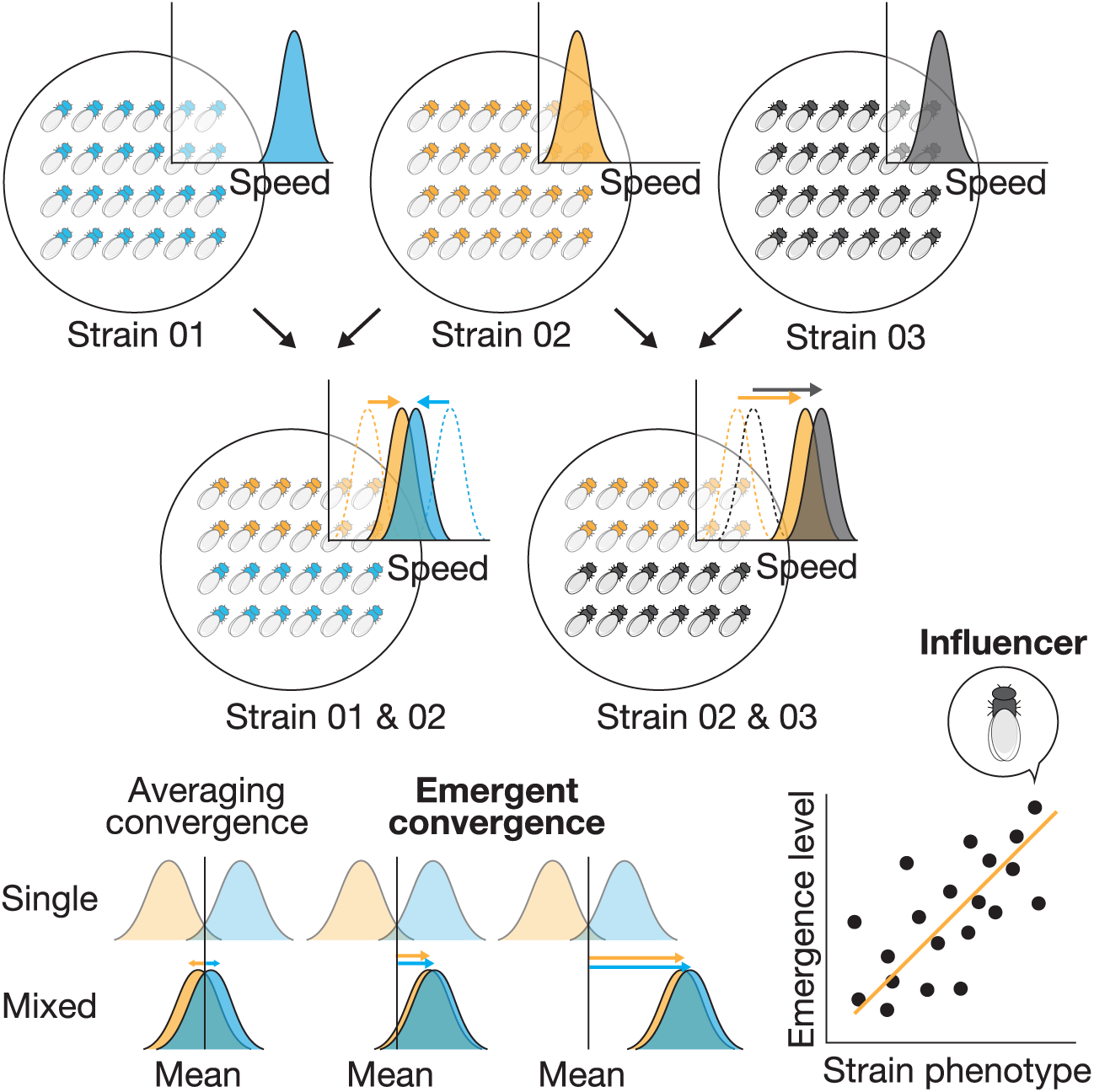
Experimental procedure. Methods for investigating the relationship between genotypic composition and group-level characteristics, and the system for detecting influential individuals.

**Figure 2.**
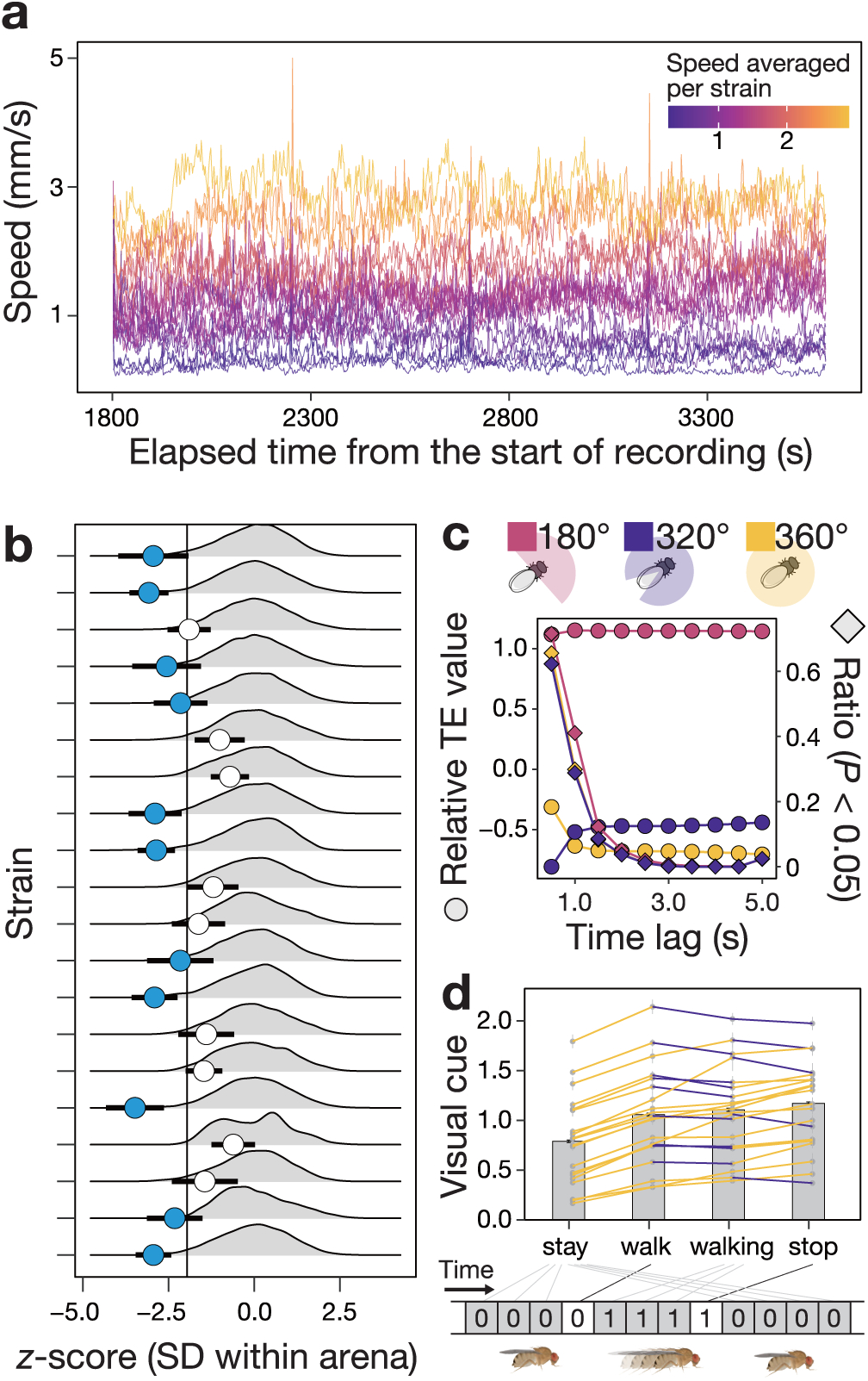
Behavioral analyses in single-strain groups. (a) The average locomotive speed of each strain at each time point. Colors indicate the average locomotive speed computed both across individuals and over time for each strain, with each line representing a specific strain. For clarity, data were averaged at 1 fps. (b) Strain-specific standardized variance within groups for locomotive speed. The gray density plot depicts the phenotypic distribution of locomotive speed in the hypothesized groups, while the points and error bars indicate the observed group means and standard errors. Blue indicates values below the vertical line (−1.96), while white indicates values above. (c) TE analysis in each field of view. Round dots indicate the amount of transferred information, while square dots indicate the corresponding p-value. (d) Visual cues present during or immediately preceding each behavioral event. Each dot with error bar shows the mean and standard error for each strain. Colors indicate yellow when values increased for the next event and blue when they decreased. The bar graph and error bar show the overall means and standard errors.

To clarify the relationship between genotypic composition and group-level characteristics, we observed 24 individuals in a group formed by mixing each strain in a 1:1 ratio (Fig. 1). Differences in mean locomotive speed between strains in a group were significantly smaller in mixed-strain groups than those calculated based on mean locomotive speed in groups consisting of a single strain (*df* = 28, *t* = 4.00, *P* < 0.001), indicating that flies conformed to the locomotive speed of others (Fig. 3a). It is noteworthy that, in a few strain combinations, the differences during mixing were larger than in other combinations. In addition, interestingly, locomotive speed in mixed-strain groups did not always converge to the arithmetic mean locomotive speed of two strains focused (i.e., the expected value under mutual conformity) (Fig. 3b). In some combinations, individuals of one strain maintained their locomotive speed, while those of the other strain unilaterally conformed to the other’s behavior. In other combinations, one strain exhibited behavior opposite to conformity, resulting in similar locomotive speeds between the two. In order to examine how strains with specific behavioral traits exhibited non-conformity, a simple linear regression analysis was conducted using the deviation from the expected locomotive speed value as the response variable, and each behavioral trait in single-strain groups as explanatory variables. The coefficient of determination (*R*^2^) varied across behavioral traits and *R*^2^ values were higher for traits relating to sociality than for other behavioral traits (Fig. 3c). The value corresponding to the moment when the sum of retinal sizes of other individuals, adjusted for inter-individual distance, reached its maximum (max sum interactions), which showed the highest *R*^2^ value, exhibited a strong correlation between single- and mixed-strain groups (Fig. 3d), indicating that “max sum interactions” remained consistent between both group types. Conversely, the position coordinates of the nearest individual at the tip of the focal fly’s “nose” (i.e., the front part of its head facing forward, position (N)), which showed higher *R*^2^ in “positioning”, were not consistent between single- and mixed-strain groups (Fig. 3e). We evaluated the influence of traits of the opposite strain on the degree of deviation in locomotive speed for each strain, using the mean behavioral traits of the opposite strain in single-strain groups as explanatory variables. The results showed that the *R*^2^ values for the opposite strain were significantly lower than those for the phenotypes of own strain, as shown in Fig. 3c (*df* = 50, *t* = 4.72, *P* < 0.001; Fig. 3f). For the focal strain trait with higher *R*^2^ (i.e., max sum interactions), *R*^2^ dropped to less than one-quarter of its original value when explained by the traits of the opposite strain.

**Figure 3.**
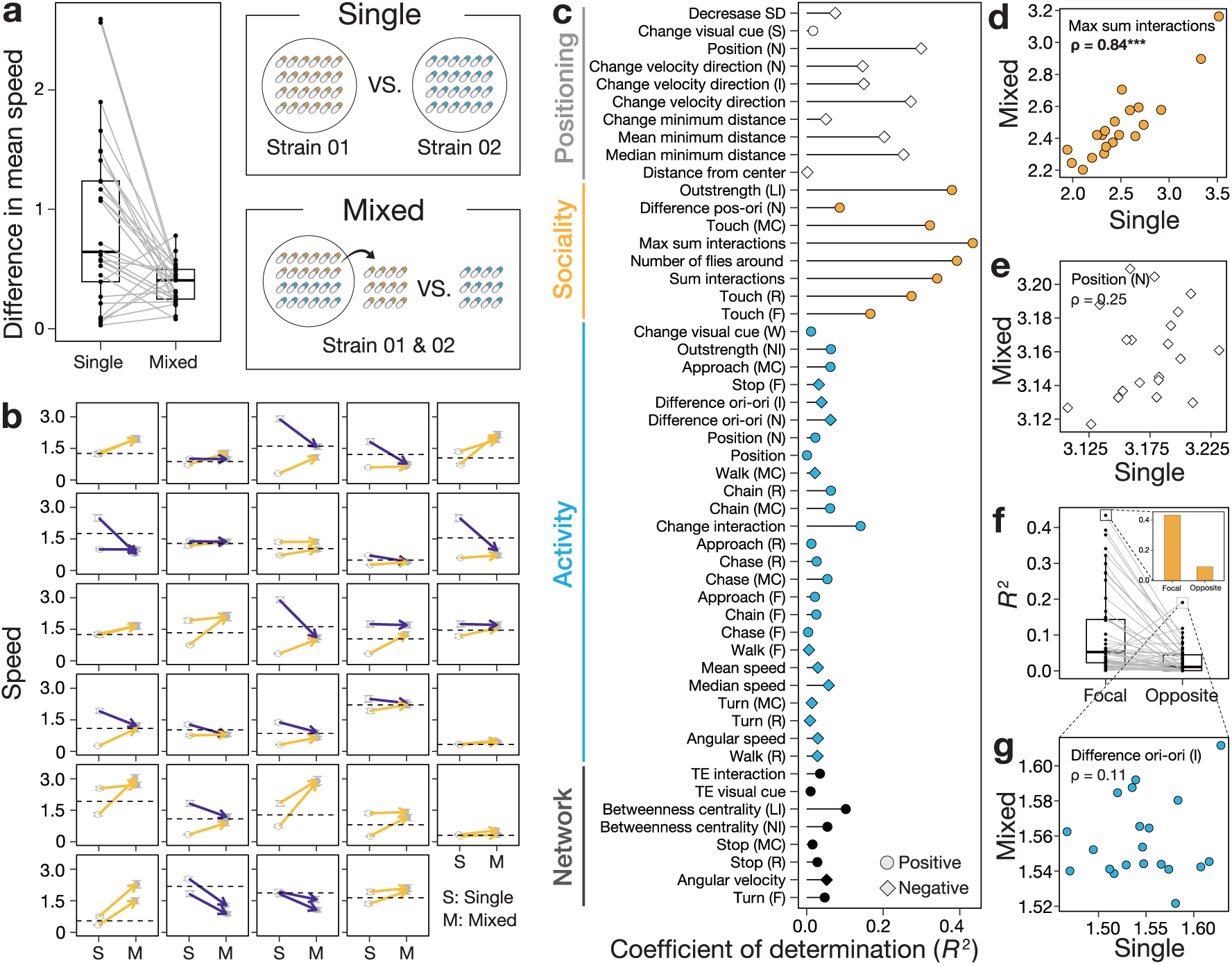
Behavioral analyses in mixed-strain groups. (a) Difference between expected and observed group locomotive speeds. Each point represents a strain combination. (b) Change in locomotive speed between single-strain and mixed-strain groups. S denotes single-strain groups, M denotes mixed-strain groups. Points and error bars indicate the mean and standard error per strain. Yellow arrows indicate increased locomotive speed from single- to mixed-strain groups, while blue arrows indicate decreased speed. Dashed lines show expected values when strains mutually conform. (c) Results of a simple linear regression model using the focal strain’s phenotype as the explanatory variable and the deviation from the expected locomotive speed as the response variable. Point colors are assigned based on clustering results. Point shapes indicate circles for positive regression slopes and squares for negative slopes. Vertical axis symbols denote: S, moment of stopping; N, nearest individual from nose tip; I, individual with largest area of projection onto the retina; LI, length of interaction; MC, maximum consistent; R, ratio; F, frequency; W, moment of walking; NI, number of interactions. Note that “pos-ori” denotes position–orientation, and “ori-ori” denotes orientation–orientation. (d) Robustness of “max sum interactions” from single- to mixed-strain conditions. ρ indicates Spearman’s correlation coefficient (\*\*\**P* < 0.001). (e) Robustness of “position (N)” from single- to mixed-strain conditions (*P* = 0.29). (f) *R*² values in simple linear models between the degree of emergent increase in locomotive speed and each trait o the focal and opposite strains. Each point represents a trait. The bar graph shows changes in the trait with the highest *R*² (max sum interactions). (g) Robustness of “difference in ori-ori” from single- to mixed-strain conditions (*P* = 0.64).

Furthermore, for the trait of the opposite strain with higher *R*^2^ (the angular difference between one’s body center and the nearest individual on the horizontal plane: difference ori-ori), *R*^2^ was 0.2, and no significant correlation was detected between single- and mixed-strain groups (Fig. 3g).

## Discussion

We demonstrated how conformity and genetic composition within groups jointly shaped the phenotype of individuals and thus group-level characteristics in *D. lutescens*. Typically, when two strains with different locomotive speeds were mixed, the locomotive speed of all individuals converged approximately to the arithmetic mean of the two strains. This result indicates that simple conformity occurred. Interestingly, an emergent increase in activity, i.e., non-additive changes in locomotive activity, was observed in groups that contained specific strains. Individuals of these strains did not conform to other strains or actively deviated from the phenotypes of others. Instead, they frequently interacted with others, which indicates high sociality. In other words, they could function as “influencers”, altering the activity levels of group members.

Indeed, flies tend to stay in areas with high visual cues. High sociality strains exhibited this tendency more strongly, potentially determining group activity levels by exerting a stronger influence than the visual cues perceived by individuals of other strains.

Numerous previous studies have demonstrated that sociality in *Drosophila* has a genetic basis (35, 36). Crucially, the theory that, for conformity-related traits, group-level characteristics and intergroup variation are shaped by the genotypic composition within the group has now been empirically verified, suggesting that a deeper understanding of highly social genotypes may be necessary to explain the behavioral variation observed in the wild.

Activity is one of the behavioral traits most strongly determined by genetics, and the heritability (*H*^2^) of activity in single-strain groups (26.3%) in the present study was similar to values reported in previous studies (37). However, even in single-strain groups, variation among groups exceeded that among strains (37.1%). Such intergroup variation is likely shaped by a combination of stochastic fluctuations and conformity. In general, the stochastic component arises from within-strain individual variation caused by factors that cannot be fully controlled, such as plasticity induced by subtle environmental differences, residual genetic variation, minor age differences, or developmental noise (38, 39). In addition, the small within-group variation observed among individuals of the same strain suggests that locomotive speed was aligned through social interactions. Therefore, even in single-strain experiments, individual variation within a strain likely produced slight intergroup variation, while phenotypes converged within each group, leading to differences in locomotive speed across groups despite the use of genetically identical individuals. However, in some strains, the within-group variation in actual groups remained similar to that in virtual groups. These strains may not only exhibit a lower degree of conformity but also less within-strain individual variation, possibly due to weaker influences of the aforementioned stochastic and plastic factors than in other strains (Fig. S1).

Previous studies have shown flies conform their walking behavior to that of others, as indicated by sensor-based activity recordings and measurements of recovery speed from freezing (40–42). The present study further elucidated the behavioral rules underlying how flies walk during social interactions, by quantitatively evaluating the immediacy of their responses using the time lag of transfer entropy (TE) and assessing the influence of the visual field. Regarding response immediacy, the time lag showing statistically clear (*P* < 0.05) causality between locomotive speed and visual cues was 0.5 seconds, accounting for approximately 70% of cases. The proportion of cases decreased to about 40% at 1 second and fell below 10% beyond 2 seconds, suggesting that flies rapidly adjust their behavior in response to the movements of other individuals.

Furthermore, at a time lag of 0.5 seconds, the proportion of individuals showing significant TE values differed by approximately 10% across visual fields, being highest at 180°. Similarly, the standardized TE value peaked at 180°, consistent with previous findings that the fly’s effective field of view is approximately 180°, and suggesting that flies are most strongly influenced by visual stimuli in the frontal visual field (43). The significant increase in visual cues immediately before walking suggests that flies were influenced to conform the movements of other individuals. Interestingly, visual cues also increased immediately before stopping, indicating non-conforming behavior.

However, previous studies have reported the opposite pattern, where individuals freeze in response to others freezing (41, 42). This discrepancy may stem from the tendency of *Drosophila* to approach and stop near stationary individuals. As a result, while it is common during free-roaming behavior for nearby individuals to begin walking, instances in which surrounding individuals stop locomotive are comparatively rare.

Therefore, the increase in visual cues observed before stopping in this study likely reflects a tendency to halt in areas with high fly density or perceived motion intensity. Indeed, moment-to-moment changes in visual cues during walking exhibited a significant positive bias, whereas no significant bias was observed during stopping, suggesting that momentary changes in visual input primarily drive walking initiation rather than termination (Fig. S2). Flies are highly social and noted for their aggregative behavior (29, 30, 44). As demonstrated here, deciphering the behavioral rules governing such moment-to-moment adjustments provides a foundation for developing mechanistic models explaining the emergence of aggregation in *Drosophila*. (45).

In mixed-strain groups, the difference in locomotive speed between strains was smaller than that in single-strain groups, indicating that conformity in locomotion occurred not only within but also across strains. However, the locomotive speed of individuals in mixed-strain groups did not always converge to the arithmetic mean of the two constituent strains. In groups containing a high-sociality strain, the locomotive speed converged to values higher than the mean, suggesting that socially interactive individuals disproportionately influenced others’ phenotypes toward their own.

Interestingly, in the mixed-strain group of the two strains showing the highest values of “max sum interactions”, the difference in locomotive speed increased from 0.076 to 0.41 (Table S1), suggesting that these two strains are specifically non-conforming to individuals of other strains. Indeed, although results from single-strain groups clearly showed conformity across all strains, it remains unclear whether this pattern is maintained in mixed-strain groups, because the difference in locomotive speed between strains within a group decreases even when one strain conforms. Previous studies have reported that *Drosophila* strains with markedly different behavioral traits fail to synchronize their activity rhythms (40).

While our hypothesis explains the mechanism in groups where one strain adjusts its locomotive speed to match the other, it does not apply to groups in which both strains increase their locomotive speeds, suggesting a possible mechanism involving the misperception of locomotive speed due to socially interactive individuals. *Drosophila* recognize another individual’s locomotive speed based on two factors: the actual locomotive speed and the inter-individual distance, a visual cue. In other words, when comparing individuals at the same locomotive speed, the closer individual is perceived to move faster. Therefore, high-sociality individuals are likely perceived as having a higher locomotive speed than their actual speed simply by approaching others, implying that strains interacting with high-sociality strains misperceive their locomotive speed and consequently increase their own locomotive speed. As a result, the high-sociality strain may conform to the elevated locomotive speed of the other strain through feedback, ultimately leading the entire group to exhibit a locomotive speed higher than expected. Furthermore, other traits showing high *R*^2^ values for deviations from expected locomotive speed were not consistent across strain mixtures, suggesting these traits are not genetically determined but rather vary in conjunction with other traits. For example, the positional relationship with other individuals tends to be determined randomly when the closest individual changes frequently. Such traits may be interactively determined by sociality and locomotive speed, and may not be functionally relevant to deviations from the expected locomotive speed.

In the present study, all strains were mixed at equal frequencies. An important question for future studies is how highly social individuals affect the phenotypes of group members when they are in the minority, as in the wild. In addition, to theoretically predict such effects, it is also important to clarify the process by which high-sociality strains shape group-level characteristics, which remains unclear.

Although the process was discussed above, these hypotheses have not been directly verified, because random simultaneous interactions between the two mixed strains make direct observation of reciprocal feedback impossible. Indeed, in the present study, several uncontrollable factors exist that may have led to emergent behavioral changes.

As discussed in the results of the single-strain groups, it cannot be ruled out that variation among individuals within a strain contributed the emergent increase in activity levels. Nevertheless, the presence of a specific strain consistently triggered emergent increase in activity levels, and the implication that certain individuals can influence the group-level characteristics represents a valuable finding. In recent years, alongside keystone species—those that play crucial roles in ecosystems (e.g., food webs)—the concept of keystone individuals (46–48), which perform similarly vital functions, have garnered attention. The socially interactive individuals identified in the present study may therefore be regarded as keystone individuals within their species, given their strong influence on the phenotypes of many conspecifics.

The present study focused on intraspecific diversity, examining the relationship between genotypic composition and group-level characteristics. A key finding is that group-level characteristics are not simply determined additively by the behavioral traits of each strain, but rather fluctuate significantly depending on the presence of specific strains. Importantly, in species like this one, which form groups composed of non-moving, randomly assembled individuals, such individuals may play a keystone role in generating intergroup variation within the population. Creating diversity that would otherwise be lost through conformity contributes to the productivity and stability of the population as a whole, extending beyond its effects on group members (49). Therefore, detecting such individuals and evaluating their cascading effects on groups may provide new perspectives for elucidating behavioral variation and predicting population dynamics in the wild.

## Materials and Methods

### Study species and rearing

*Drosophila* flies are well known for exhibiting strong sociality and converging many behavioral traits, including locomotive speed, through social interactions (40, 41). We used *D. lutescens*, which belongs to the *melanogaster* group and is easily maintained under laboratory conditions. The observation systems used for *D. melanogaster* can also be applied to this species. The iso-female strains used here were established in 2020 (38), indicating that they were more recently collected from the wild than most *D. melanogaster* strains maintained at stock centers, suggesting that these strains have not undergone selection under laboratory conditions and thus still retain wild-type variation. We used 20 iso-female strains (Table S2) of this species maintained in an incubator under constant conditions (22℃, 12L12D). The flies were reared in plastic vials (30 mm in diameter and 100 mm in height) on a nutritive medium according to the protocol of Fitzpatrick et al. (50) to obtain individuals for experiments. The flies used in all experiments were offspring obtained from parent flies that had been allowed to lay eggs for one week. The offspring were transferred to new vials immediately after emergence and aged for 1–7 days before use.

### Video recording

Flies were observed in an open-field test using a circular arena (140 mm in diameter and 2.5 mm in height) with a shallow concave shape similar to that of a mortar (Fig. S3) (51). Female flies were randomly selected from a vial, anesthetized with CO₂, and transferred to the arena. Female individuals were used because they exhibit higher social activity than males (44), and male–male interactions such as aggression and same-sex courtship could confound behavioral measurements (52, 53). After the flies were introduced into the arena, they were placed in an incubator set at 25°C, and their behavior was recorded from above using a video camera (Fig. 4). Video recording lasted for 1 hour; the first 30 minutes served as an acclimation period, and the remaining 30 minutes were used for behavioral analysis.

**Figure 4.**
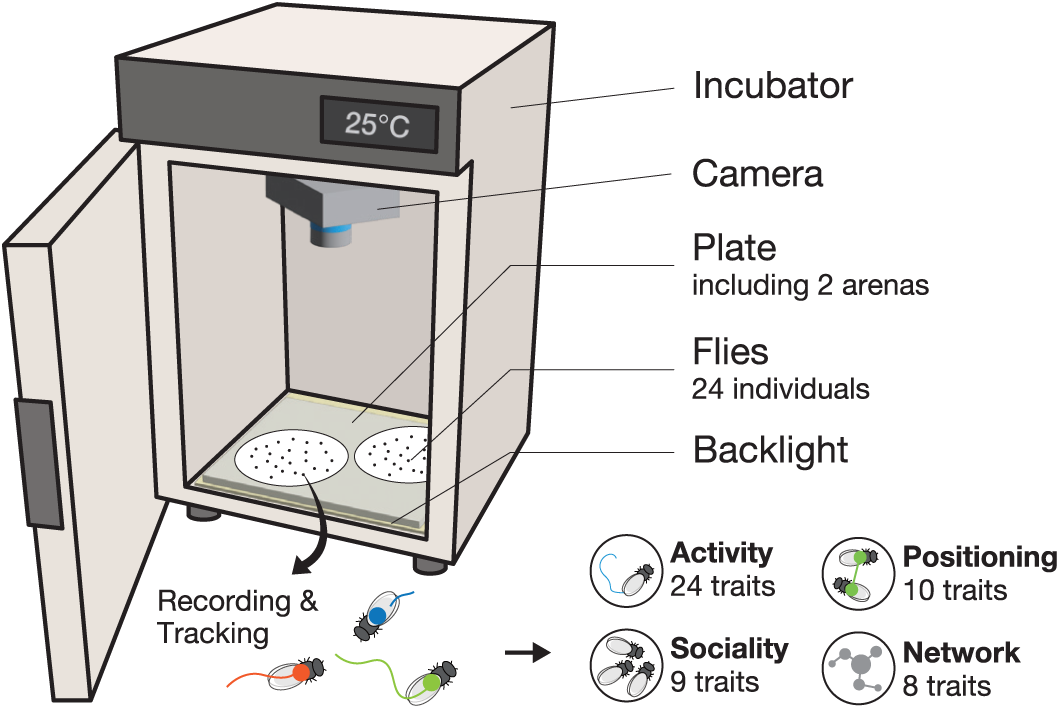
Experimental equipment. All recordings were conducted in a controlled environment within incubators. Each recording session used one plate containing two arenas, each holding 24 flies.

Recordings were performed for 29 strain combinations, with each strain used in at least two different group compositions. In the single-strain condition, anesthetized individuals were placed at the center of the arena at the start of the video recording. In the two-strain condition, 12 individuals from each strain were randomly selected from separate vials, and their initial positions were set at two distinct locations along the y-axis from the center, allowing strain identification during tracking. To standardize the initial spatial configuration between the two setups, the starting positions were placed closer to the center to minimize spatial dispersion.

### Quantification of behavioral traits

We obtained X–Y coordinates and orientations of each individual in the videos using FlyTracker (34). The videos used for tracking were recorded at 6 frames per second (fps); however, behavioral traits were quantified from tracking data resampled at 2 fps. We quantified locomotor speed and the visual cue (see below for details) as primary behavioral traits from the tracking data. Furthermore, a total of 51 behavioral traits (Table S3) (31), derived from individual coordinate and orientation data, were quantified and hierarchically clustered using Ward’s method into 4 categories: activity (25 traits), sociality (8 traits), positioning (10 traits), and network (8 traits). The clusters broadly represent traits associated with locomotive behavior (activity), social interactions and aggregation (sociality), spatial relationships between specific pairs of individuals (positioning), and network structure (network). Since the number of clusters was chosen visually, these categories were not incorporated into the statistical analyses and were used solely as descriptive labels to aid interpretation (Fig. S4). The visual cues were calculated as in Ferreira and Moita et al. (41) as an index that evaluates the degree of motion of other individuals as reflected on the retina, because *Drosophila* flies use this cue to conform to the walking and stopping of others. A visual cue can be written as the following equation:

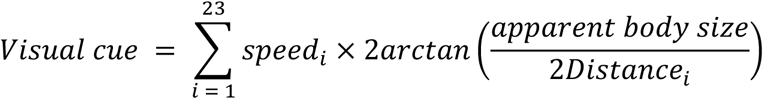

where *i* is an individual in the arena, and “*apparent body size*” represents the projected size on the retina, approximated by an ellipse with a major axis of 3 mm and a minor axis of 1.5 mm. The visual cue as an index was considered the effects of the flies’ visual field, because both the distance between individuals and their visual field were important for quantifying networks and interactions. The visual field was defined as 180°, 320°, and 360° in the calculation of all behavioral traits (Fig. S5). To indicate that the visual cue was used for flies’ walking and stopping, the causality between the flies’ locomotive speed and the visual cue was calculated using transfer entropy (TE) (54, 55). TE for two observed time series *x*_t_ and *y*_t_ can be written as the following equation:

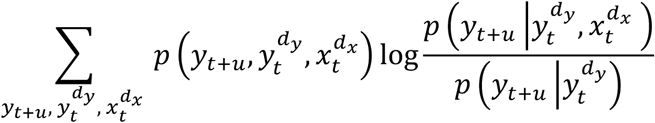

where *t* is a discrete-valued time index and *u* denotes the prediction time, a discrete-valued time interval. *y*_t_^dy^ and *x*_t_^dx^ are *d*_x_ - and *d*_y_-dimensional delay vectors.

### Verification of Conformity

If individuals within a group exhibited conformity, the variation among interacting individuals would be expected to be smaller than that among non-interacting individuals. Even among individuals of the same strain, behavioral fluctuations arise from genetic diversity and developmental noise (56). Nevertheless, and inter-individual variation among those that interact within a group is expected to decrease through phenotypic convergence, even within a single strain. Therefore, to test whether conformity occurred, we constructed virtual groups from the empirical data and calculated the phenotypic variation among non-interacting individuals. For each strain, 10,000 trials were performed by randomly sampling 24 individuals regardless of their group membership, and the extent of phenotypic variation among non-interacting individuals was computed. We compared within-group variation in virtual groups constructed without interaction structure with that in actual groups.

### Assessment of changes in the final convergent locomotive speed

To investigate the mechanisms underlying the final convergent locomotive speed, we analyzed how strain-specific behavioral patterns relate to changes of locomotive speed. However, mixed-strain groups are expected to fall into two categories: groups in which members conform and move at an average speed, and groups in which the presence of strain-specific behavioral patterns causes their locomotive speed to deviate from this average. Therefore, only the overall locomotive speed of mixed-strain groups, or the simple change in speed of each strain relative to its behavioral patterns, cannot accurately identify the factors causing deviations from the average. We first calculated the average locomotive speed of single-strain groups and used this conformity-based average as the expected value. We then computed the deviation from that value using the following equation:

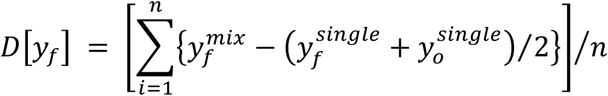

where *y*_f_ and *y*_o_are the locomotive speeds of the focal and opposite strains, respectively; “*single*” and “*mix*” indicate single-strain and mixed-strain groups; and *n* represents the number of opposite strains paired with each focal strain (i.e., strain combinations).

### Statistical analyses

All statistical analyses were performed in R (version 4.3.0). The R package *R.matlab* was used to import coordinate data obtained from video tracking. Behavioral differences were primarily analyzed using linear mixed-effects models (LMMs) implemented in the *lmerTest* package. Experimental condition was included as a fixed effect, while random effects were specified according to the data structure. For the LMM analyzed via ANOVA with the *car* package, group identity was included as a random effect to account for variability between groups. The contribution of each factor to variation in locomotive speed was estimated using the *VCA* package, based on the same LMM structure used for the ANOVA. Locomotive speed was log-transformed prior to analysis. For the LMM analyzed using *t*-tests, strain, the strain × group interaction, and the strain × group × individual interaction were included as random effects to account for repeated measures. Models were fitted using restricted maximum likelihood (REML). Simple linear models (LMs) were used to analyze the effects of continuous predictors on outcomes when no random effects were needed. Model assumptions of normality and homoscedasticity were assessed visually. Relationships between variables were assessed using correlation analyses (three correlation tests in total), and paired *t*-tests (two tests in total) were used to compare conditions.

### Ethics statement

No ethical approval was required for the present study, as it involved non-invasive observation of an invertebrate species that is neither endangered nor protected. All individuals were collected within the facilities of our institution and did not require specific permissions.

## Acknowledgments

This research was supported by the Chiba University Comprehensive and Challenging Fusion Innovator Doctoral Program, by the Japan Society for the Promotion of Science (Grants-in-Aid for Scientific Research 22H05646 and 23H03840 to Y.T.), and by the Sasakawa Research Grant (2023-5051 to K.H.).

