## Supplemental Information for "Influencer in flies: Socially interactive individuals shape group-level characteristics"

### Supporting Information for
### Influencer in flies: Socially interactive individuals shape group-level characteristics

Kaiya Hamamichi[1] and Yuma Takahashi[2]

1. Graduate School of Science and Engineering, Chiba University, Chiba, Japan
2. Graduate School of Science, Chiba University, Chiba, Japan

**This PDF file includes:**

Supporting text

Figures S1 to S5

Tables S1 to S3

SI References

**Dataset in Figshare (separate file).**

- df_boot_share.csv
- time_series_share.csv
- traits_data_share.csv

### Supporting Information Text

#### For Results

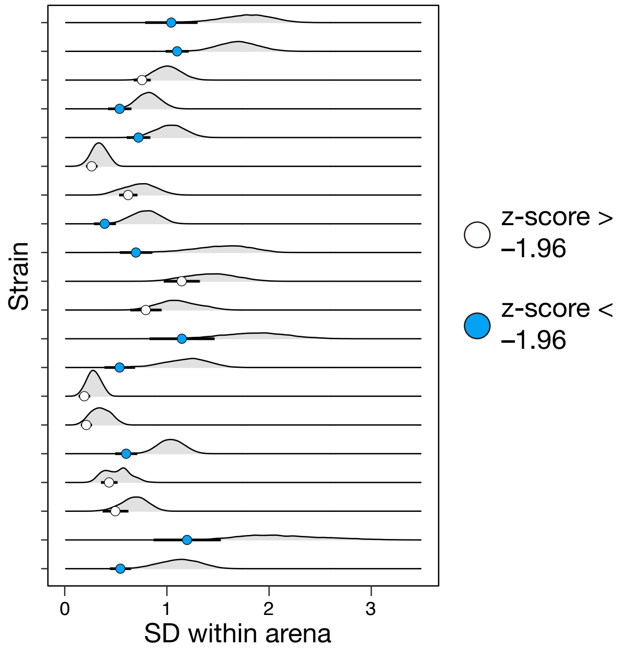


**Figure S1.** Within-group variance in locomotive speed of each strain prior to standardization.

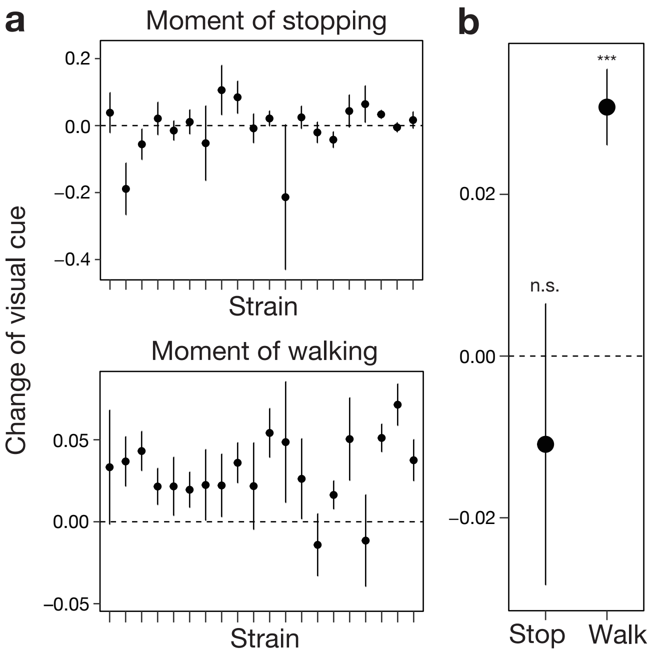


**Figure S2.** Moment-to-moment changes in the visual cues immediately before walking and stopping. Results for each strain (a), and the overall averaged result (b). The statistical tests were performed using a *t*-test with a linear mixed model (LMM) that included random effects for strain and the interactions between strain × group and strain × group × individual (***: $P < 0.001$, N.S.: $P > 0.05$).

### Table

**S1 table**. Sociality (max sum interactions) of each strain and differences in locomotive speed when mixed with other strains.

| line1 | line2 | Sociality in line1 | Sociality in line2 | Mean of sociality | Difference in single | Difference in mix |
|---|---|---|---|---|---|---|
| L054 | L077 | 2.917 | 2.436 | 2.676 | 0.643 | 0.424 |
| L054 | L355 | 2.917 | 2.480 | 2.698 | 0.566 | 0.248 |
| L054 | L104 | 2.917 | 2.326 | 2.621 | 1.091 | 0.446 |
| L054 | L508 | 2.917 | 2.683 | 2.800 | 0.617 | 0.266 |
| L059 | L355 | 2.305 | 2.480 | 2.393 | 0.573 | 0.496 |
| L059 | L083 | 2.305 | 2.251 | 2.278 | 1.898 | 0.267 |
| L059 | L539 | 2.305 | 2.201 | 2.253 | 1.489 | 0.198 |
| L077 | L104 | 2.436 | 2.326 | 2.381 | 0.448 | 0.080 |
| L083 | L591 | 2.251 | 2.335 | 2.293 | 1.232 | 0.303 |
| L085 | L387 | 2.510 | 3.515 | 3.013 | 0.053 | 0.485 |
| L085 | L510 | 2.510 | 2.593 | 2.551 | 0.526 | 0.201 |
| L099 | L581 | 1.990 | 2.346 | 2.168 | 2.564 | 0.457 |
| L099 | L135 | 1.990 | 2.415 | 2.202 | 1.409 | 0.515 |
| L099 | L591 | 1.990 | 2.335 | 2.162 | 1.484 | 0.310 |
| L099 | L104 | 1.990 | 2.326 | 2.158 | 0.081 | 0.205 |
| L099 | L508 | 1.990 | 2.683 | 2.336 | 0.393 | 0.779 |
| L099 | L535 | 1.990 | 1.944 | 1.967 | 0.029 | 0.104 |

| | | | | | | |
|---|---|---|---|---|---|---|
| L104 | L355 | 2.326 | 2.480 | 2.403 | 1.657 | 0.574 |
| L122 | L135 | 2.649 | 2.415 | 2.532 | 0.579 | 0.240 |
| L122 | L137 | 2.649 | 2.735 | 2.692 | 0.212 | 0.227 |
| L137 | L535 | 2.735 | 1.944 | 2.340 | 1.071 | 0.273 |
| L355 | L510 | 2.480 | 2.593 | 2.537 | 1.166 | 0.539 |
| L355 | L591 | 2.480 | 2.335 | 2.408 | 0.093 | 0.651 |
| L387 | L538 | 3.515 | 3.330 | 3.423 | 0.076 | 0.412 |
| L508 | L591 | 2.683 | 2.335 | 2.509 | 1.091 | 0.511 |
| L508 | L539 | 2.683 | 2.201 | 2.442 | 0.267 | 0.314 |
| L535 | L581 | 1.944 | 2.346 | 2.145 | 2.593 | 0.508 |
| L538 | L544 | 3.330 | 2.105 | 2.717 | 1.236 | 0.411 |
| L544 | L591 | 2.105 | 2.335 | 2.220 | 0.712 | 0.405 |

38

#### For Materials and Methods

**The threshold in quantified traits.** Several traits quantified here (such as the number of walking bouts) were quantified using arbitrary thresholds. The choice of thresholds was generally based on prior research (1, 2).

**Behavioral consistency**: For traits converted to binary values (0 or 1), actions lasting less than one second were assigned a value of 0. After this processing, if an action was restarted within 4 seconds of its termination, these actions were treated as a single continuous sequence.

**Interaction**: Individuals were defined as interacting if they were within the field of view (180°) and remained within 2.5 times their body length (3 mm) for at least 1 second.

**Touch**: When the distance between individuals was within one body length (3 mm), they were defined as being in touching.

**Walk**: To prevent tracking coordinate drift caused by video quality from being misidentified as walking, movement was defined as walking only when the locomotive speed exceeded 0.5 mm/s.

**Turn**: A turn was defined as occurring when the body orientation change exceeded 0.01 rad/s.

**Chase**: Individuals were defined as being in a chase when:

- The difference in body orientation between the chasing and chased individuals was within 180°
- Both individuals were walking
- Both were within 2.5 body lengths of each other
- The chased individual was within the chasing individual's field of view

**Chain**: When three or more individuals were linked in a chase, they were defined as forming a chain.

**Approach**: An approaching individual was defined as approaching another when:

- The approaching individual was walking
- The approached individual was within 2.5 body lengths
- The approached individual was within the approaching individual's field of view
- The distance between the individuals was decreasing

### Tables

**S2 table**. All sample information used in the analysis.

| Date | Time | ID | Arena | Line1 | Line2 | Age |
|---|---|---|---|---|---|---|
| 22-Jul-22 | 13:07 | 20220722_01 | a01 | | L538 | 3_6 |
| 22-Jul-22 | 13:07 | 20220722_01 | a02 | | L539 | 3_6 |
| 22-Jul-22 | 13:07 | 20220722_DJI_01 | a01 | | L508 | 3_6 |
| 22-Jul-22 | 13:07 | 20220722_DJI_01 | a02 | | L387 | 3_6 |
| 22-Jul-22 | 14:25 | 20220722_DJI_02 | a01 | | L538 | 3_6 |
| 22-Jul-22 | 15:50 | 20220722_02 | a01 | L387 | L538 | 3_6 |
| 22-Jul-22 | 15:50 | 20220722_02 | a02 | | L538 | 3_6 |
| 22-Jul-22 | 15:50 | 20220722_DJI_03 | a01 | | L508 | 3_6 |
| 27-Jul-22 | 14:42 | 20220727_01 | a02 | | L355 | 1_4 |
| 27-Jul-22 | 16:10 | 20220727_02 | a02 | L508 | L539 | 1_4 |
| 8-Aug-22 | 13:56 | 20220808_01 | a01 | | L508 | 3_6 |
| 8-Aug-22 | 13:56 | 20220808_01 | a02 | | L508 | 3_6 |
| 8-Aug-22 | 13:44 | 20220808_DJI_01 | a01 | | L510 | 3_6 |
| 8-Aug-22 | 13:44 | 20220808_DJI_01 | a02 | | L510 | 3_6 |
| 8-Aug-22 | 15:15 | 20220808_DJI_02 | a01 | | L539 | 3_6 |
| 8-Aug-22 | 15:15 | 20220808_DJI_02 | a02 | | L539 | 3_6 |
| 8-Aug-22 | 16:33 | 20220808_03 | a01 | | L538 | 3_6 |
| 8-Aug-22 | 16:33 | 20220808_03 | a02 | | L538 | 3_6 |
| 8-Aug-22 | 16:19 | 20220808_DJI_03 | a01 | | L387 | 3_6 |

| | | | | | | |
|---|---|---|---|---|---|---|
| 8-Aug-22 | 16:19 | 20220808_DJI_03 | a02 | | L387 | 3_6 |
| 9-Aug-22 | 12:56 | 20220809_01 | a01 | | L355 | 4_7 |
| 9-Aug-22 | 12:56 | 20220809_01 | a02 | | L355 | 4_7 |
| 9-Aug-22 | 13:07 | 20220809_DJI_01 | a01 | | L591 | 4_7 |
| 9-Aug-22 | 13:07 | 20220809_DJI_01 | a02 | | L591 | 4_7 |
| 9-Aug-22 | 14:13 | 20220809_02 | a01 | L535 | L581 | 4_7 |
| 9-Aug-22 | 14:13 | 20220809_02 | a02 | | L535 | 4_7 |
| 9-Aug-22 | 14:28 | 20220809_DJI_02 | a01 | | L054 | 4_7 |
| 9-Aug-22 | 14:28 | 20220809_DJI_02 | a02 | | L054 | 4_7 |
| 9-Aug-22 | 15:38 | 20220809_03 | a01 | | L083 | 4_7 |
| 9-Aug-22 | 15:38 | 20220809_03 | a02 | | L083 | 4_7 |
| 9-Aug-22 | 15:49 | 20220809_DJI_03 | a01 | | L077 | 4_7 |
| 9-Aug-22 | 15:49 | 20220809_DJI_03 | a02 | | L077 | 4_7 |
| 9-Aug-22 | 16:45 | 20220809_04 | a01 | | L122 | 4_7 |
| 9-Aug-22 | 16:45 | 20220809_04 | a02 | | L122 | 4_7 |
| 9-Aug-22 | 16:57 | 20220809_DJI_04 | a01 | | L099 | 4_7 |
| 9-Aug-22 | 16:57 | 20220809_DJI_04 | a02 | | L099 | 4_7 |
| 12-Aug-22 | 13:14 | 20220812_01 | a01 | | L510 | 1_3 |
| 12-Aug-22 | 13:14 | 20220812_01 | a02 | | L510 | 1_3 |
| 12-Aug-22 | 14:18 | 20220812_02 | a01 | | L387 | 1_3 |
| 12-Aug-22 | 14:18 | 20220812_02 | a02 | | L387 | 1_3 |
| 12-Aug-22 | 14:29 | 20220812_DJI_02 | a01 | L508 | L054 | 1_3 |
| 12-Aug-22 | 14:29 | 20220812_DJI_02 | a02 | | L508 | 1_3 |

| | | | | | | |
|---|---|---|---|---|---|---|
| 12-Aug-22 | 15:20 | 20220812_03 | a01 | L591 | L083 | 1_3 |
| 12-Aug-22 | 15:20 | 20220812_03 | a02 | | L591 | 1_3 |
| 12-Aug-22 | 15:35 | 20220812_DJI_03 | a01 | L539 | L059 | 1_3 |
| 12-Aug-22 | 15:35 | 20220812_DJI_03 | a02 | | L539 | 1_3 |
| 12-Aug-22 | 16:26 | 20220812_04 | a01 | | L077 | 1_3 |
| 12-Aug-22 | 16:26 | 20220812_04 | a02 | | L077 | 1_3 |
| 12-Aug-22 | 16:41 | 20220812_DJI_04 | a01 | | L059 | 1_3 |
| 12-Aug-22 | 16:41 | 20220812_DJI_04 | a02 | | L059 | 1_3 |
| 14-Aug-22 | 12:29 | 20220814_01 | a01 | | L099 | 3_5 |
| 14-Aug-22 | 12:29 | 20220814_01 | a02 | | L099 | 3_5 |
| 14-Aug-22 | 12:36 | 20220814_DJI_01 | a01 | | L085 | 3_5 |
| 14-Aug-22 | 12:36 | 20220814_DJI_01 | a02 | | L085 | 3_5 |
| 14-Aug-22 | 13:45 | 20220814_02 | a01 | | L137 | 3_5 |
| 14-Aug-22 | 13:45 | 20220814_02 | a02 | L122 | L137 | 3_5 |
| 14-Aug-22 | 13:58 | 20220814_DJI_02 | a01 | L104 | L077 | 3_5 |
| 14-Aug-22 | 13:58 | 20220814_DJI_02 | a02 | | L135 | 3_5 |
| 14-Aug-22 | 14:49 | 20220814_03 | a01 | | L077 | 3_5 |
| 14-Aug-22 | 14:49 | 20220814_03 | a02 | | L077 | 3_5 |
| 14-Aug-22 | 14:59 | 20220814_DJI_03 | a01 | | L387 | 3_5 |
| 14-Aug-22 | 14:59 | 20220814_DJI_03 | a02 | | L387 | 3_5 |
| 14-Aug-22 | 16:03 | 20220814_04 | a01 | L077 | L054 | 3_5 |
| 14-Aug-22 | 16:03 | 20220814_04 | a02 | | L099 | 3_5 |
| 14-Aug-22 | 16:17 | 20220814_DJI_04 | a01 | L059 | L083 | 3_5 |

| | | | | | | |
|---|---|---|---|---|---|---|
| 17-Aug-22 | 12:49 | 20220817_01 | a01 | | L104 | 2_5 |
| 17-Aug-22 | 12:49 | 20220817_01 | a02 | | L104 | 2_5 |
| 17-Aug-22 | 12:59 | 20220817_DJI_01 | a01 | | L059 | 2_5 |
| 17-Aug-22 | 12:59 | 20220817_DJI_01 | a02 | | L059 | 2_5 |
| 17-Aug-22 | 13:50 | 20220817_02 | a01 | | L538 | 2_5 |
| 17-Aug-22 | 13:50 | 20220817_02 | a02 | | L538 | 2_5 |
| 17-Aug-22 | 14:51 | 20220817_03 | a01 | | L137 | 2_5 |
| 17-Aug-22 | 14:51 | 20220817_03 | a02 | | L137 | 2_5 |
| 17-Aug-22 | 15:01 | 20220817_DJI_03 | a01 | | L099 | 2_5 |
| 17-Aug-22 | 15:01 | 20220817_DJI_03 | a02 | | L099 | 2_5 |
| 17-Aug-22 | 15:52 | 20220817_04 | a01 | | L054 | 2_5 |
| 17-Aug-22 | 15:52 | 20220817_04 | a02 | | L054 | 2_5 |
| 17-Aug-22 | 16:02 | 20220817_DJI_04 | a01 | | L122 | 2_5 |
| 17-Aug-22 | 16:02 | 20220817_DJI_04 | a02 | | L122 | 2_5 |
| 18-Aug-22 | 13:29 | 20220818_01 | a01 | L085 | L387 | 3_6 |
| 18-Aug-22 | 13:29 | 20220818_01 | a02 | | L085 | 3_6 |
| 18-Aug-22 | 13:37 | 20220818_DJI_01 | a01 | | L054 | 3_6 |
| 18-Aug-22 | 13:37 | 20220818_DJI_01 | a02 | | L054 | 3_6 |
| 18-Aug-22 | 14:35 | 20220818_02 | a01 | L510 | L355 | 3_6 |
| 18-Aug-22 | 14:35 | 20220818_02 | a02 | | L510 | 3_6 |
| 18-Aug-22 | 14:46 | 20220818_DJI_02 | a01 | L083 | L591 | 3_6 |
| 18-Aug-22 | 14:46 | 20220818_DJI_02 | a02 | | L083 | 3_6 |
| 18-Aug-22 | 15:36 | 20220818_03 | a01 | | L508 | 3_6 |

| | | | | | | |
|---|---|---|---|---|---|---|
| 18-Aug-22 | 15:36 | 20220818_03 | a02 | | L355 | 3_6 |
| 18-Aug-22 | 15:47 | 20220818_DJI_03 | a01 | L077 | L104 | 3_6 |
| 18-Aug-22 | 15:47 | 20220818_DJI_03 | a02 | | L077 | 3_6 |
| 18-Aug-22 | 16:37 | 20220818_04 | a01 | L581 | L099 | 3_6 |
| 18-Aug-22 | 16:37 | 20220818_04 | a02 | | L539 | 3_6 |
| 18-Aug-22 | 16:50 | 20220818_DJI_04 | a01 | | L104 | 3_6 |
| 18-Aug-22 | 16:50 | 20220818_DJI_04 | a02 | | L591 | 3_6 |
| 19-Aug-22 | 13:22 | 20220819_01 | a01 | L077 | L054 | 1_3 |
| 19-Aug-22 | 13:22 | 20220819_01 | a02 | L077 | L054 | 1_3 |
| 19-Aug-22 | 13:33 | 20220819_DJI_01 | a01 | L077 | L054 | 1_3 |
| 19-Aug-22 | 13:33 | 20220819_DJI_01 | a02 | | L059 | 1_3 |
| 19-Aug-22 | 14:34 | 20220819_02 | a01 | | L387 | 1_3 |
| 19-Aug-22 | 14:34 | 20220819_02 | a02 | | L387 | 1_3 |
| 19-Aug-22 | 14:42 | 20220819_DJI_02 | a01 | | L510 | 1_3 |
| 19-Aug-22 | 14:42 | 20220819_DJI_02 | a02 | | L510 | 1_3 |
| 19-Aug-22 | 15:37 | 20220819_03 | a01 | L508 | L539 | 1_3 |
| 19-Aug-22 | 15:37 | 20220819_03 | a02 | | L508 | 1_3 |
| 19-Aug-22 | 15:46 | 20220819_DJI_03 | a01 | L137 | L535 | 1_3 |
| 19-Aug-22 | 15:46 | 20220819_DJI_03 | a02 | | L137 | 1_3 |
| 19-Aug-22 | 16:38 | 20220819_04 | a01 | | L085 | 1_3 |
| 19-Aug-22 | 16:38 | 20220819_04 | a02 | | L085 | 1_3 |
| 19-Aug-22 | 16:49 | 20220819_DJI_04 | a01 | L083 | L591 | 1_3 |
| 19-Aug-22 | 16:49 | 20220819_DJI_04 | a02 | | L083 | 1_3 |

| | | | | | | |
|---|---|---|---|---|---|---|
| 22-Aug-22 | 15:13 | 20220822_01 | a02 | L099 | L135 | 4_6 |
| 22-Aug-22 | 15:30 | 20220822_DJI_01 | a01 | L538 | L387 | 4_6 |
| 22-Aug-22 | 15:30 | 20220822_DJI_01 | a02 | L538 | L387 | 4_6 |
| 23-Aug-22 | 12:52 | 20220823_01 | a01 | L054 | L077 | 1_4 |
| 23-Aug-22 | 12:52 | 20220823_01 | a02 | | L054 | 1_4 |
| 23-Aug-22 | 13:06 | 20220823_DJI_01 | a01 | L077 | L054 | 1_4 |
| 23-Aug-22 | 13:06 | 20220823_DJI_01 | a02 | L077 | L104 | 1_4 |
| 23-Aug-22 | 13:58 | 20220823_02 | a01 | L137 | L122 | 1_4 |
| 23-Aug-22 | 13:58 | 20220823_02 | a02 | | L137 | 1_4 |
| 23-Aug-22 | 14:08 | 20220823_DJI_02 | a01 | L104 | L355 | 1_4 |
| 23-Aug-22 | 14:08 | 20220823_DJI_02 | a02 | L104 | L355 | 1_4 |
| 23-Aug-22 | 15:07 | 20220823_03 | a01 | L135 | L122 | 1_4 |
| 23-Aug-22 | 15:07 | 20220823_03 | a02 | | L137 | 1_4 |
| 23-Aug-22 | 15:19 | 20220823_DJI_03 | a01 | L054 | L508 | 1_4 |
| 23-Aug-22 | 15:19 | 20220823_DJI_03 | a02 | L054 | L508 | 1_4 |
| 24-Aug-22 | 15:48 | 20220824_DJI_01 | a01 | L387 | L085 | 1_5 |
| 24-Aug-22 | 15:48 | 20220824_DJI_01 | a02 | L387 | L085 | 1_5 |
| 25-Aug-22 | 13:30 | 20220825_DJI_01 | a01 | L083 | L591 | 3_6 |
| 25-Aug-22 | 13:30 | 20220825_DJI_01 | a02 | L083 | L591 | 3_6 |
| 26-Aug-22 | 13:27 | 20220826_DJI_01 | a01 | L059 | L083 | 1_3 |
| 26-Aug-22 | 13:27 | 20220826_DJI_01 | a02 | L059 | L083 | 1_3 |
| 26-Aug-22 | 13:38 | 20220826_DJI2_01 | a01 | L054 | L508 | 1_3 |
| 26-Aug-22 | 13:38 | 20220826_DJI2_01 | a02 | L054 | L508 | 1_3 |

| | | | | | | |
|---|---|---|---|---|---|---|
| 26-Aug-22 | 15:35 | 20220826_DJI_03 | a01 | L387 | L085 | 1_3 |
| 26-Aug-22 | 15:35 | 20220826_DJI_03 | a02 | L387 | L085 | 1_3 |
| 26-Aug-22 | 15:46 | 20220826_DJI2_03 | a01 | L099 | L135 | 1_3 |
| 26-Aug-22 | 15:46 | 20220826_DJI2_03 | a02 | L099 | L135 | 1_3 |
| 26-Aug-22 | 16:37 | 20220826_DJI_04 | a01 | L122 | L137 | 1_3 |
| 26-Aug-22 | 16:37 | 20220826_DJI_04 | a02 | L122 | L137 | 1_3 |
| 26-Aug-22 | 16:50 | 20220826_DJI2_04 | a01 | L355 | L104 | 1_3 |
| 26-Aug-22 | 16:50 | 20220826_DJI2_04 | a02 | L535 | L581 | 1_3 |
| 27-Aug-22 | 13:39 | 20220827_DJI_01 | a01 | L539 | L508 | 2_4 |
| 27-Aug-22 | 13:39 | 20220827_DJI_01 | a02 | L539 | L508 | 2_4 |
| 27-Aug-22 | 13:52 | 20220827_DJI2_01 | a01 | L355 | L510 | 2_4 |
| 27-Aug-22 | 13:52 | 20220827_DJI2_01 | a02 | L085 | L510 | 2_4 |
| 27-Aug-22 | 14:41 | 20220827_DJI_02 | a01 | L591 | L083 | 2_4 |
| 27-Aug-22 | 14:41 | 20220827_DJI_02 | a02 | L591 | L083 | 2_4 |
| 27-Aug-22 | 14:53 | 20220827_DJI2_02 | a01 | L387 | L538 | 2_4 |
| 27-Aug-22 | 14:53 | 20220827_DJI2_02 | a02 | L387 | L538 | 2_4 |
| 27-Aug-22 | 15:43 | 20220827_DJI_03 | a01 | L122 | L137 | 2_4 |
| 27-Aug-22 | 15:43 | 20220827_DJI_03 | a02 | L059 | L083 | 2_4 |
| 27-Aug-22 | 15:56 | 20220827_DJI2_03 | a01 | L135 | L099 | 2_4 |
| 27-Aug-22 | 15:56 | 20220827_DJI2_03 | a02 | L135 | L099 | 2_4 |
| 27-Aug-22 | 16:48 | 20220827_DJI_04 | a01 | L099 | L135 | 2_4 |
| 27-Aug-22 | 16:48 | 20220827_DJI_04 | a02 | L122 | L135 | 2_4 |
| 27-Aug-22 | 17:01 | 20220827_DJI2_04 | a01 | | L539 | 2_4 |

| | | | | | | |
|---|---|---|---|---|---|---|
| 27-Aug-22 | 17:01 | 20220827_DJI2_04 | a02 | | L539 | 2_4 |
| 30-Aug-22 | 13:04 | 20220830_DJI_01 | a01 | L135 | L099 | 1_4 |
| 30-Aug-22 | 13:04 | 20220830_DJI_01 | a02 | | L135 | 1_4 |
| 30-Aug-22 | 13:15 | 20220830_DJI2_01 | a01 | L510 | L085 | 1_4 |
| 30-Aug-22 | 13:15 | 20220830_DJI2_01 | a02 | | L581 | 1_4 |
| 30-Aug-22 | 14:25 | 20220830_DJI_02 | a01 | L054 | L077 | 1_7 |
| 30-Aug-22 | 14:25 | 20220830_DJI_02 | a02 | L104 | L077 | 1_7 |
| 30-Aug-22 | 14:38 | 20220830_DJI2_02 | a01 | L104 | L077 | 1_7 |
| 30-Aug-22 | 14:38 | 20220830_DJI2_02 | a02 | L104 | L077 | 1_7 |
| 30-Aug-22 | 15:32 | 20220830_DJI_03 | a01 | L387 | L538 | 1_7 |
| 30-Aug-22 | 15:43 | 20220830_DJI2_03 | a01 | L137 | L122 | 1_7 |
| 30-Aug-22 | 15:43 | 20220830_DJI2_03 | a02 | | L137 | 1_4 |
| 30-Aug-22 | 16:33 | 20220830_DJI_04 | a01 | L510 | L355 | 1_7 |
| 30-Aug-22 | 16:33 | 20220830_DJI_04 | a02 | | L355 | 1_4 |
| 30-Aug-22 | 16:46 | 20220830_DJI2_04 | a02 | | L591 | 1_4 |
| 31-Aug-22 | 13:34 | 20220831_DJI_01 | a01 | L508 | L539 | 2_5 |
| 31-Aug-22 | 13:34 | 20220831_DJI_01 | a02 | L508 | L539 | 2_5 |
| 31-Aug-22 | 13:46 | 20220831_DJI2_01 | a01 | L135 | L122 | 2_5 |
| 31-Aug-22 | 13:46 | 20220831_DJI2_01 | a02 | L135 | L122 | 2_5 |
| 31-Aug-22 | 14:43 | 20220831_DJI_02 | a01 | L137 | L535 | 2_5 |
| 31-Aug-22 | 14:43 | 20220831_DJI_02 | a02 | L538 | L387 | 2_5 |
| 31-Aug-22 | 14:55 | 20220831_DJI2_02 | a01 | L083 | L059 | 2_5 |
| 31-Aug-22 | 14:55 | 20220831_DJI2_02 | a02 | L083 | L059 | 2_5 |

| | | | | | | |
|---|---|---|---|---|---|---|
| 31-Aug-22 | 15:58 | 20220831_DJI2_03 | a01 | L355 | L104 | 2_5 |
| 31-Aug-22 | 15:58 | 20220831_DJI2_03 | a02 | | L104 | 2_5 |
| 31-Aug-22 | 16:46 | 20220831_DJI_04 | a01 | L099 | L581 | 2_5 |
| 31-Aug-22 | 16:46 | 20220831_DJI_04 | a02 | L510 | L085 | 2_5 |
| 31-Aug-22 | 16:58 | 20220831_DJI2_04 | a01 | L122 | L137 | 2_5 |
| 31-Aug-22 | 16:58 | 20220831_DJI2_04 | a02 | | L122 | 2_5 |
| 1-Sep-22 | 13:19 | 20220901_DJI_01 | a02 | L077 | L104 | 3_6 |
| 1-Sep-22 | 13:29 | 20220901_DJI2_01 | a01 | L059 | L539 | 3_6 |
| 1-Sep-22 | 13:29 | 20220901_DJI2_01 | a02 | | L059 | 3_6 |
| 1-Sep-22 | 14:26 | 20220901_DJI_02 | a01 | L508 | L054 | 3_6 |
| 1-Sep-22 | 14:26 | 20220901_DJI_02 | a02 | L508 | L054 | 3_6 |
| 1-Sep-22 | 14:37 | 20220901_DJI2_02 | a01 | L077 | L104 | 3_6 |
| 1-Sep-22 | 14:37 | 20220901_DJI2_02 | a02 | | L122 | 3_6 |
| 1-Sep-22 | 15:26 | 20220901_DJI_03 | a02 | | L104 | 3_6 |
| 1-Sep-22 | 15:40 | 20220901_DJI2_03 | a01 | L083 | L059 | 3_6 |
| 1-Sep-22 | 15:40 | 20220901_DJI2_03 | a02 | L135 | L122 | 3_6 |
| 1-Sep-22 | 16:32 | 20220901_DJI_04 | a01 | | L085 | 3_6 |
| 1-Sep-22 | 16:32 | 20220901_DJI_04 | a02 | | L083 | 3_6 |
| 2-Sep-22 | 13:17 | 20220902_DJI_01 | a01 | L355 | L104 | 1_7 |
| 2-Sep-22 | 13:17 | 20220902_DJI_01 | a02 | L535 | L137 | 1_7 |
| 2-Sep-22 | 13:30 | 20220902_DJI2_01 | a01 | L539 | L059 | 1_3 |
| 2-Sep-22 | 13:30 | 20220902_DJI2_01 | a02 | L539 | L059 | 1_3 |
| 2-Sep-22 | 14:27 | 20220902_DJI_02 | a01 | L085 | L510 | 1_3 |

| | | | | | | |
|---|---|---|---|---|---|---|
| 2-Sep-22 | 14:27 | 20220902_DJI_02 | a02 | L085 | L510 | 1_3 |
| 2-Sep-22 | 14:40 | 20220902_DJI2_02 | a01 | | L083 | 1_3 |
| 2-Sep-22 | 14:40 | 20220902_DJI2_02 | a02 | | L135 | 1_3 |
| 2-Sep-22 | 15:42 | 20220902_DJI_03 | a02 | | L591 | 1_3 |
| 2-Sep-22 | 15:57 | 20220902_DJI2_03 | a01 | L122 | L135 | 1_3 |
| 2-Sep-22 | 15:57 | 20220902_DJI2_03 | a02 | L122 | L135 | 1_3 |
| 2-Sep-22 | 16:54 | 20220902_DJI_04 | a01 | | L059 | 1_3 |
| 2-Sep-22 | 16:54 | 20220902_DJI_04 | a02 | | L591 | 1_3 |
| 2-Sep-22 | 17:10 | 20220902_DJI2_04 | a01 | L135 | L122 | 1_3 |
| 2-Sep-22 | 17:10 | 20220902_DJI2_04 | a02 | L539 | L508 | 1_3 |
| 6-Sep-22 | 12:53 | 20220906_DJI_01 | a01 | L387 | L085 | 5_7 |
| 6-Sep-22 | 12:53 | 20220906_DJI_01 | a02 | L510 | L085 | 5_7 |
| 6-Sep-22 | 13:07 | 20220906_DJI2_01 | a01 | L539 | L059 | 5_7 |
| 6-Sep-22 | 13:07 | 20220906_DJI2_01 | a02 | | L083 | 1_4 |
| 6-Sep-22 | 14:07 | 20220906_DJI_02 | a01 | L539 | L059 | 1_4 |
| 6-Sep-22 | 14:07 | 20220906_DJI_02 | a02 | L539 | L059 | 1_4 |
| 6-Sep-22 | 14:19 | 20220906_DJI2_02 | a01 | L137 | L535 | 1_4 |
| 6-Sep-22 | 14:19 | 20220906_DJI2_02 | a02 | | L104 | 1_4 |
| 6-Sep-22 | 15:18 | 20220906_DJI_03 | a01 | L581 | L099 | 1_4 |
| 6-Sep-22 | 15:18 | 20220906_DJI_03 | a02 | L581 | L099 | 1_4 |
| 6-Sep-22 | 15:29 | 20220906_DJI2_03 | a01 | L104 | L355 | 1_4 |
| 6-Sep-22 | 15:29 | 20220906_DJI2_03 | a02 | | L104 | 1_4 |
| 6-Sep-22 | 16:51 | 20220906_DJI_04 | a01 | L510 | L085 | 1_4 |

| | | | | | | |
|---|---|---|---|---|---|---|
| 6-Sep-22 | 16:51 | 20220906_DJI_04 | a02 | | L085 | 1_4 |
| 6-Sep-22 | 17:03 | 20220906_DJI2_04 | a01 | L387 | L085 | 1_4 |
| 6-Sep-22 | 17:03 | 20220906_DJI2_04 | a02 | | L355 | 1_4 |
| 7-Sep-22 | 14:26 | 20220907_DJI_01 | a01 | L510 | L355 | 2_5 |
| 7-Sep-22 | 14:26 | 20220907_DJI_01 | a02 | L104 | L355 | 2_5 |
| 7-Sep-22 | 14:37 | 20220907_DJI2_01 | a01 | L581 | L099 | 2_5 |
| 7-Sep-22 | 14:37 | 20220907_DJI2_01 | a02 | | L122 | 2_5 |
| 7-Oct-22 | 12:58 | 20221007_DJI_01 | a01 | L535 | L137 | 1_7 |
| 7-Oct-22 | 12:58 | 20221007_DJI_01 | a02 | | L535 | 4_7 |
| 7-Oct-22 | 13:11 | 20221007_DJI2_01 | a01 | L581 | L535 | 1_7 |
| 7-Oct-22 | 13:11 | 20221007_DJI2_01 | a02 | | L581 | 4_7 |
| 7-Oct-22 | 14:10 | 20221007_DJI_02 | a01 | L510 | L355 | 1_7 |
| 7-Oct-22 | 14:10 | 20221007_DJI_02 | a02 | | L135 | 4_7 |
| 7-Oct-22 | 14:21 | 20221007_DJI2_02 | a01 | | L535 | 1_7 |
| 7-Oct-22 | 14:21 | 20221007_DJI2_02 | a02 | | L135 | 1_3 |
| 14-Oct-22 | 15:40 | 20221014_DJI_01 | a01 | L535 | L581 | 1_7 |
| 14-Oct-22 | 15:40 | 20221014_DJI_01 | a02 | | L581 | 1_7 |
| 14-Oct-22 | 15:46 | 20221014_DJI2_01 | a01 | | L535 | 1_7 |
| 11-Nov-22 | 15:06 | 20221111_DJI2_01 | a01 | | L581 | 3_6 |
| 11-Nov-22 | 15:06 | 20221111_DJI2_01 | a02 | | L581 | 3_6 |
| 8-Feb-23 | 16:45 | 20230208_DJI2_04 | a01 | L059 | L355 | 5_7 |
| 8-Feb-23 | 16:45 | 20230208_DJI2_04 | a02 | L059 | L355 | 5_7 |
| 13-Feb-23 | 14:53 | 20230213_DJI_01 | a01 | L535 | L099 | 3_5 |

| | | | | | | |
|---|---|---|---|---|---|---|
| 13-Feb-23 | 14:53 | 20230213_DJI_01 | a02 | L535 | L099 | 3_5 |
| 15-Feb-23 | 13:14 | 20230215_DJI_01 | a01 | L535 | L099 | 1_7 |
| 15-Feb-23 | 13:14 | 20230215_DJI_01 | a02 | L535 | L099 | 1_7 |
| 15-Feb-23 | 14:40 | 20230215_DJI_02 | a01 | L059 | L355 | 5_7 |
| 15-Feb-23 | 14:40 | 20230215_DJI_02 | a02 | L059 | L355 | 5_7 |
| 15-Feb-23 | 16:46 | 20230215_DJI_04 | a02 | L355 | L059 | 5_7 |
| 7-Sep-23 | 16:15 | 20230907_DJI2_02 | a02 | L137 | L581 | 3_6 |
| 26-Jun-24 | 14:20 | 20240626_DJI_01 | a01 | L054 | L077 | 5_6 |
| 26-Jun-24 | 14:20 | 20240626_DJI_01 | a02 | | L099 | 5_6 |
| 26-Jun-24 | 14:46 | 20240626_DJI2_01 | a01 | L591 | L083 | 1_4 |
| 26-Jun-24 | 14:46 | 20240626_DJI2_01 | a02 | | L591 | 5_6 |
| 26-Jun-24 | 15:28 | 20240626_DJI_02 | a01 | | L054 | 1_4 |
| 26-Jun-24 | 15:28 | 20240626_DJI_02 | a02 | L054 | L077 | 1_4 |
| 26-Jun-24 | 15:49 | 20240626_DJI2_02 | a01 | | L355 | 1_4 |
| 26-Jun-24 | 15:49 | 20240626_DJI2_02 | a02 | | L059 | 1_4 |
| 26-Jun-24 | 16:32 | 20240626_DJI_03 | a01 | | L122 | 1_4 |
| 26-Jun-24 | 16:32 | 20240626_DJI_03 | a02 | | L122 | 1_4 |
| 26-Jun-24 | 16:52 | 20240626_DJI2_03 | a01 | | L099 | 1_4 |
| 26-Jun-24 | 16:52 | 20240626_DJI2_03 | a02 | L535 | L099 | 1_4 |
| 27-Jun-24 | 14:37 | 20240627_DJI_01 | a01 | | L104 | 2_5 |
| 27-Jun-24 | 14:37 | 20240627_DJI_01 | a02 | L104 | L355 | 2_5 |
| 27-Jun-24 | 14:51 | 20240627_DJI2_01 | a01 | | L077 | 2_5 |
| 27-Jun-24 | 14:51 | 20240627_DJI2_01 | a02 | L355 | L510 | 2_5 |

| | | | | | | |
|---|---|---|---|---|---|---|
| 27-Jun-24 | 16:36 | 20240627_DJI_02 | a01 | L355 | L059 | 1_7 |
| 27-Jun-24 | 16:36 | 20240627_DJI_02 | a02 | L355 | L510 | 1_7 |
| 1-Jul-24 | 13:42 | 20240701_DJI_01 | a01 | L059 | L355 | 3_5 |
| 3-Jul-24 | 13:43 | 20240703_DJI_01 | a01 | | L059 | 1_4 |
| 3-Jul-24 | 13:43 | 20240703_DJI_01 | a02 | L099 | L581 | 1_4 |
| 3-Jul-24 | 13:53 | 20240703_DJI2_01 | a01 | | L059 | 1_4 |
| 3-Jul-24 | 13:53 | 20240703_DJI2_01 | a02 | | L059 | 1_4 |
| 3-Jul-24 | 14:45 | 20240703_DJI_02 | a01 | | L137 | 1_4 |
| 3-Jul-24 | 14:45 | 20240703_DJI_02 | a02 | | L083 | 1_4 |
| 8-Jul-24 | 16:42 | 20240708_DJI_01 | a01 | | L581 | 3_5 |
| 8-Jul-24 | 16:42 | 20240708_DJI_01 | a02 | L137 | L535 | 3_5 |
| 10-Jul-24 | 13:36 | 20240710_DJI_01 | a01 | | L535 | 1_4 |
| 10-Jul-24 | 13:36 | 20240710_DJI_01 | a02 | L581 | L099 | 1_4 |
| 10-Jul-24 | 13:49 | 20240710_DJI2_01 | a01 | | L535 | 1_4 |
| 10-Jul-24 | 13:49 | 20240710_DJI2_01 | a02 | L538 | L544 | 1_4 |
| 10-Jul-24 | 14:45 | 20240710_DJI_02 | a01 | | L137 | 1_4 |
| 10-Jul-24 | 14:45 | 20240710_DJI_02 | a02 | | L535 | 1_4 |
| 12-Jul-24 | 16:32 | 20240712_DJI_01 | a02 | L099 | L591 | 3_6 |
| 16-Jul-24 | 13:12 | 20240716_DJI_01 | a01 | L535 | L581 | 4_6 |
| 16-Jul-24 | 13:12 | 20240716_DJI_01 | a02 | L535 | L137 | 4_6 |
| 16-Jul-24 | 13:22 | 20240716_DJI2_01 | a02 | L544 | L538 | 4_6 |
| 16-Jul-24 | 14:49 | 20240716_DJI2_02 | a02 | L591 | L508 | 4_6 |
| 16-Jul-24 | 15:41 | 20240716_DJI_03 | a02 | L591 | L508 | 4_6 |

| | | | | | | |
|---|---|---|---|---|---|---|
| 16-Jul-24 | 15:54 | 20240716_DJI2_03 | a01 | L104 | L054 | 4_6 |
| 16-Jul-24 | 15:54 | 20240716_DJI2_03 | a02 | L104 | L054 | 4_6 |
| 17-Jul-24 | 13:26 | 20240717_DJI_01 | a01 | | L581 | 1_4 |
| 17-Jul-24 | 13:26 | 20240717_DJI_01 | a02 | | L544 | 1_4 |
| 17-Jul-24 | 13:38 | 20240717_DJI2_01 | a01 | L535 | L137 | 1_4 |
| 17-Jul-24 | 13:38 | 20240717_DJI2_01 | a02 | L535 | L137 | 1_4 |
| 17-Jul-24 | 14:38 | 20240717_DJI_02 | a02 | L544 | L538 | 1_4 |
| 17-Jul-24 | 14:49 | 20240717_DJI2_02 | a02 | L099 | L535 | 1_4 |
| 17-Jul-24 | 15:47 | 20240717_DJI_03 | a01 | L544 | L538 | 1_4 |
| 17-Jul-24 | 15:47 | 20240717_DJI_03 | a02 | L544 | L538 | 1_4 |
| 18-Jul-24 | 15:05 | 20240718_DJI_02 | a02 | L104 | L054 | 2_5 |
| 18-Jul-24 | 15:16 | 20240718_DJI2_02 | a01 | L508 | L591 | 2_5 |
| 18-Jul-24 | 15:16 | 20240718_DJI2_02 | a02 | L508 | L591 | 2_5 |
| 22-Jul-24 | 13:26 | 20240722_DJI_01 | a01 | L535 | L581 | 3_5 |
| 22-Jul-24 | 13:26 | 20240722_DJI_01 | a02 | L535 | L581 | 3_5 |
| 22-Jul-24 | 13:39 | 20240722_DJI2_01 | a01 | L099 | L104 | 3_5 |
| 22-Jul-24 | 13:39 | 20240722_DJI2_01 | a02 | L099 | L104 | 3_5 |
| 22-Jul-24 | 14:43 | 20240722_DJI_02 | a02 | L508 | L591 | 3_5 |
| 22-Jul-24 | 14:55 | 20240722_DJI2_02 | a01 | L099 | L104 | 3_5 |
| 22-Jul-24 | 14:55 | 20240722_DJI2_02 | a02 | L099 | L535 | 3_5 |
| 22-Jul-24 | 15:49 | 20240722_DJI_03 | a01 | L538 | L544 | 3_5 |
| 22-Jul-24 | 15:49 | 20240722_DJI_03 | a02 | L538 | L544 | 3_5 |
| 22-Jul-24 | 15:56 | 20240722_DJI2_03 | a01 | | L544 | 3_5 |

| | | | | | | |
|---|---|---|---|---|---|---|
| 22-Jul-24 | 16:59 | 20240722_DJI2_04 | a02 | L054 | L104 | 3_5 |
| 23-Jul-24 | 14:32 | 20240723_DJI_01 | a01 | L099 | L591 | 4_6 |
| 23-Jul-24 | 14:32 | 20240723_DJI_01 | a02 | L099 | L591 | 4_6 |
| 23-Jul-24 | 14:44 | 20240723_DJI2_01 | a01 | L099 | L104 | 4_6 |
| 23-Jul-24 | 14:44 | 20240723_DJI2_01 | a02 | L099 | L104 | 4_6 |
| 23-Jul-24 | 15:47 | 20240723_DJI_02 | a01 | L099 | L591 | 4_6 |
| 23-Jul-24 | 15:47 | 20240723_DJI_02 | a02 | L099 | L591 | 4_6 |
| 23-Jul-24 | 16:02 | 20240723_DJI2_02 | a01 | L099 | L508 | 4_6 |
| 23-Jul-24 | 16:02 | 20240723_DJI2_02 | a02 | L099 | L508 | 4_6 |
| 24-Jul-24 | 15:54 | 20240724_DJI2_03 | a01 | | L544 | 1_4 |
| 24-Jul-24 | 15:54 | 20240724_DJI2_03 | a02 | L544 | L591 | 1_4 |
| 24-Jul-24 | 16:55 | 20240724_DJI2_04 | a02 | L355 | L591 | 1_4 |
| 25-Jul-24 | 16:46 | 20240725_DJI_01 | a02 | L355 | L054 | 2_5 |
| 25-Jul-24 | 16:56 | 20240725_DJI2_01 | a01 | L508 | L099 | 2_5 |
| 25-Jul-24 | 16:56 | 20240725_DJI2_01 | a02 | L508 | L099 | 2_5 |
| 26-Jul-24 | 16:47 | 20240726_DJI_01 | a01 | L104 | L054 | 3_6 |
| 26-Jul-24 | 16:47 | 20240726_DJI_01 | a02 | L104 | L054 | 3_6 |
| 26-Jul-24 | 16:57 | 20240726_DJI2_01 | a01 | L099 | L508 | 3_6 |
| 26-Jul-24 | 16:57 | 20240726_DJI2_01 | a02 | L099 | L508 | 3_6 |
| 30-Jul-24 | 14:44 | 20240730_DJI2_02 | a02 | L104 | L054 | 4_6 |
| 30-Jul-24 | 15:40 | 20240730_DJI_03 | a01 | L508 | L591 | 4_6 |
| 30-Jul-24 | 15:40 | 20240730_DJI_03 | a02 | L508 | L591 | 4_6 |
| 30-Jul-24 | 15:53 | 20240730_DJI2_03 | a01 | L099 | L104 | 4_6 |

| | | | | | | |
|---|---|---|---|---|---|---|
| 30-Jul-24 | 15:53 | 20240730_DJI2_03 | a02 | L099 | L104 | 4_6 |
| 30-Jul-24 | 16:40 | 20240730_DJI_04 | a01 | | L544 | 4_6 |
| 30-Jul-24 | 16:40 | 20240730_DJI_04 | a02 | L508 | L099 | 4_6 |
| 30-Jul-24 | 16:54 | 20240730_DJI2_04 | a01 | L355 | L054 | 4_6 |
| 30-Jul-24 | 16:54 | 20240730_DJI2_04 | a02 | L355 | L054 | 4_6 |
| 31-Jul-24 | 13:50 | 20240731_DJI_01 | a01 | L355 | L054 | 1_7 |
| 31-Jul-24 | 13:50 | 20240731_DJI_01 | a02 | L355 | L054 | 1_7 |
| 31-Jul-24 | 15:59 | 20240731_DJI2_01 | a01 | | L544 | 1_4 |
| 31-Jul-24 | 15:59 | 20240731_DJI2_01 | a02 | L544 | L591 | 1_7 |
| 31-Jul-24 | 15:13 | 20240731_DJI_02 | a01 | | L544 | 1_4 |
| 31-Jul-24 | 15:13 | 20240731_DJI_02 | a02 | | L544 | 1_4 |
| 31-Jul-24 | 15:24 | 20240731_DJI2_02 | a01 | L099 | L591 | 1_4 |
| 31-Jul-24 | 15:24 | 20240731_DJI2_02 | a02 | L099 | L591 | 1_4 |
| 2-Aug-24 | 15:06 | 20240802_DJI_01 | a02 | L544 | L591 | 3_6 |
| 2-Aug-24 | 15:13 | 20240802_DJI2_01 | a02 | L544 | L591 | 3_6 |
| 5-Aug-24 | 13:40 | 20240805_DJI2_01 | a01 | L544 | L591 | 3_5 |
| 5-Aug-24 | 13:40 | 20240805_DJI2_01 | a02 | L544 | L591 | 3_5 |
| 5-Aug-24 | 16:09 | 20240805_DJI_03 | a01 | L544 | L591 | 3_5 |
| 5-Aug-24 | 16:09 | 20240805_DJI_03 | a02 | L544 | L591 | 3_5 |
| 21-Aug-24 | 14:49 | 20240821_DJI_01 | a01 | L355 | L054 | 4_7 |
| 21-Aug-24 | 14:49 | 20240821_DJI_01 | a02 | L355 | L054 | 4_7 |
| 23-Aug-24 | 14:00 | 20240823_DJI_01 | a01 | L355 | L591 | 2_5 |
| 23-Aug-24 | 14:00 | 20240823_DJI_01 | a02 | L355 | L591 | 2_5 |

| | | | | | | |
|---|---|---|---|---|---|---|
| 23-Aug-24 | 14:07 | 20240823_DJI2_01 | a02 | L355 | L591 | 2_5 |
| 23-Aug-24 | 16:05 | 20240823_DJI_02 | a01 | L591 | L355 | 2_5 |
| 23-Aug-24 | 16:05 | 20240823_DJI_02 | a02 | L591 | L355 | 2_5 |
| 13-Dec-24 | 15:14 | 20241213_DJI_01 | a01 | | L135 | 3_6 |
| 13-Dec-24 | 15:14 | 20241213_DJI_01 | a02 | | L135 | 3_6 |
| 20-Dec-24 | 15:48 | 20241220_DJI2_01 | a01 | | L054 | 2_6 |
| 20-Dec-24 | 15:48 | 20241220_DJI2_01 | a02 | L355 | L591 | 2_6 |

**Table S3.** Meanings of traits quantified in the present study.

| **Traits in the dataset** | **Traits in the paper** | **Description** |
|---|---|---|
| mean_sd | Decresase SD | Standard deviation of locomotive speed within a group, standardized within each strain |
| diff_visual_cue180_stop | Change visual cue (S) | Moment-to-moment change in the visual cue just before stopping |
| n_cloang_opp | Position (N) | Position of focal individual relative to the individual closest to the tip of the nose (rad) |
| n_diff_veldir | Change velocity direction (N) | Difference in velocity direction between the individual closest to the tip of the nose and the focal individual |
| visual_cue180_NS_int_diff_veldir | Change velocity direction (I) | Difference in velocity direction between the largest individual in the field of view and the focal individual |
| speed_dir_change | Change velocity direction | Average change in velocity direction per second |
| change_min_dist | Change minimum distance | Change in minimum distance between the focal individual's body center and other individual' body centers (mm/s) |

| | | |
|---|---|---|
| min_dist_mean | Mean minimum distance | Mean of minimum distance between the focal individual's body center and other individual' body centers |
| min_dist_med | Median minimum distance | Median of minimum distance between the focal individual's body center and other individual' body centers |
| dist_center | Distance from center | The distance from arena center |
| str_loi180 | Outstrength (LI) | Length of interactions of a focal individual |
| n_diff_cloang_ori | Difference pos-ori (N) | The difference between the focal individual's orientation and the position of individual closest to the tip of the nose |
| max_touch_con | Touch (MC) | Maximum consistent of touch with other individuals |
| int_max_total180 | Max sum interactions | Maximum total area of other individuals in the field of view |
| num_ind_2body | Number of flies around | Number of individuals within twice the body length |
| int_sum180 | Sum interactions | Mean of total area of other individuals in the field of view |
| touch_ratio | Touch (R) | Ratio of touch with other individuals |
| touch_fre | Touch (F) | Frequency of touch with other individuals |

| | | |
|---|---|---|
| diff_visual_cue180_walk | Change visual cue (W) | Moment-to-moment change in the visual cue just before walking |
| str_noi180 | Outstrength (NI) | Number of interactions of a certain individual |
| max_app_con | Approach (MC) | Maximum consistent of approaching other individuals |
| stop_freq | Stop (F) | Frequency of stopping |
| visual_cue180_NS_int_diff_ori | Difference ori-ori (I) | The difference in orientation between the focal individual and the largest individual in the field of view |
| n_diff_ori | Difference ori-ori (N) | The difference in orientation between the focal individual and individual closest to the tip of the nose |
| n_angle_closest | Position (N) | Position of the individual closest to the tip of the nose relative to focal individual |
| angle_closest | Position | Position of the individual closest to the body center relative to focal individual |
| max_walk_con | Walk (MC) | Maximum consistent of walking |
| chain_ratio | Chain (R) | Ratio of participation in chain |
| max_chain_con | Chain (MC) | Maximum consistent of participation in chain |
| int_change180 | Change interaction | Change in total area of other individuals in the field of view (rad/s) |
| app_ratio | Approach (R) | Ratio of approaching other individuals |
| chase_ratio | Chase (R) | Ratio of chasing other individuals |

| | | |
|---|---|---|
| max_chase_con | Chase (MC) | Maximum consistent of chasing other individuals |
| app_fre | Approach (F) | Frequency of approaching other individuals |
| chain_fre | Chain (F) | Frequency of participation in chain |
| chase_fre | Chase (F) | Frequency of chasing other individuals |
| speed_fre | Walk (F) | Frequency of walking |
| mean_speed | Mean speed | Mean of locomotive speed |
| med_speed | Median speed | Median of locomotive speed |
| max_turn_con | Turn (MC) | Maximum consistent of changes in the orientation |
| turn_ratio | Turn (R) | Ratio of changes in the orientation |
| angular_speed | Angular speed | Mean of angular speed |
| walk_ratio | Walk (R) | Ratio of walking |
| TE_conforming_visual_cue180_NS_lag1 | TE interaction | Mean degree of information transfer to the focal individual's locomotive speed from the total area of other individuals within the field of view |
| TE_conforming_visual_cue180_lag1 | TE visual cue | Mean degree of information transfer to the focal individual's locomotive speed from the visual cue |
| loi_bet180 | Betweenness centrality (LI) | Betweenness centrality calculated based on the length of interactions |
| noi_bet180 | Betweenness centrality (NI) | Betweenness centrality calculated based on the number of interactions |
| max_stop_con | Stop (MC) | Maximum consistent of stopping |
| stop_ratio | Stop (R) | Ratio of stopping |

| | | |
|---|---|---|
| ang_vel | Angular velocity | Mean of angular velocity |
| turn_fre | Turn (F) | Frequency of changes in the orientation |

### Figures

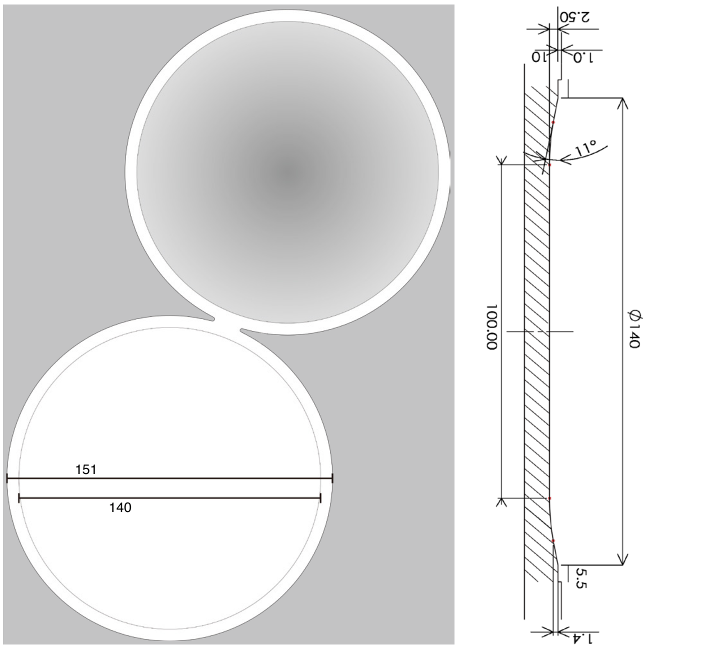


**Figure S3.** The arena blueprint used in the present study

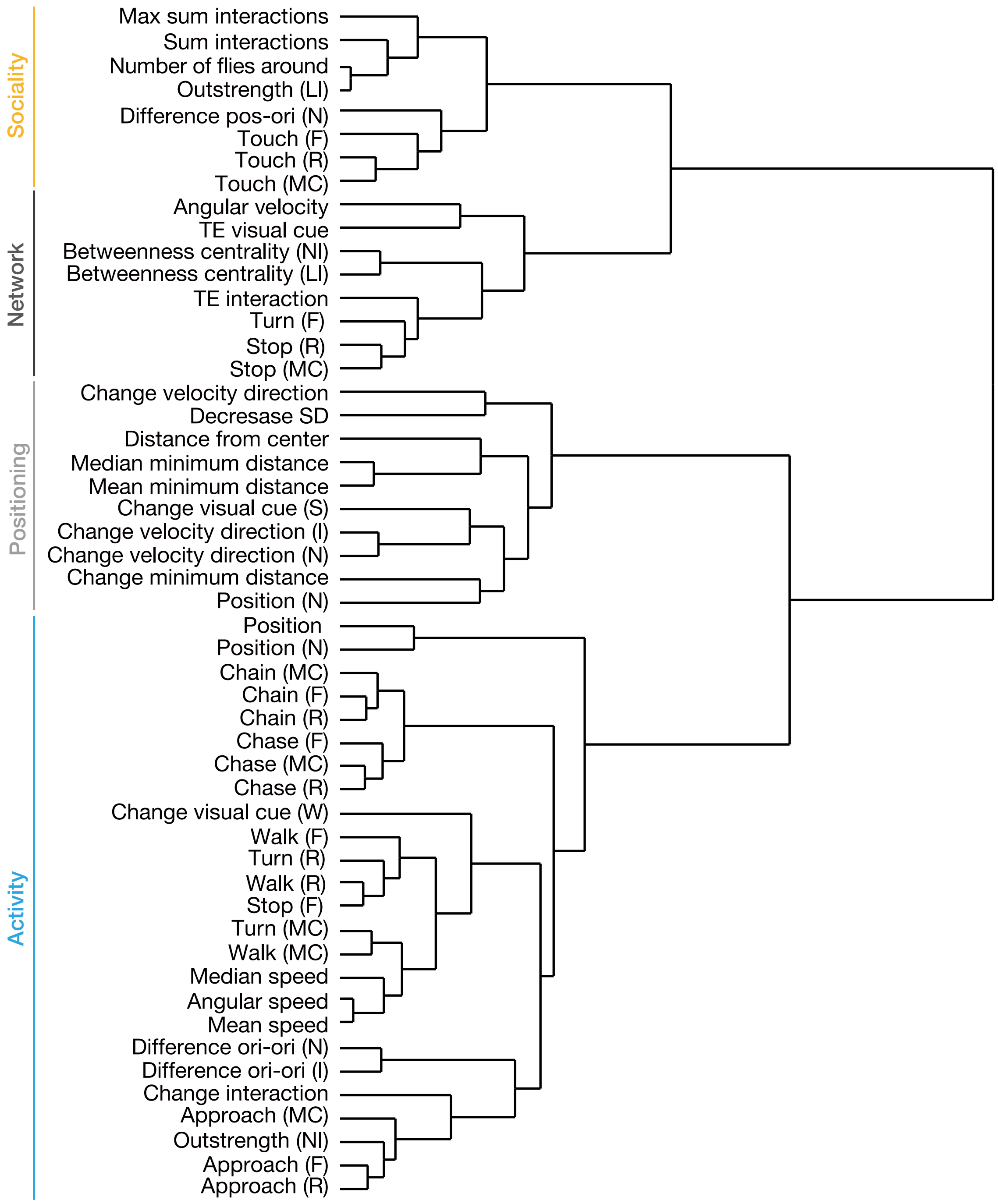


**Figure S4.** The dendrogram estimated using Ward's method based on phenotypic data from individuals in single-strain groups.

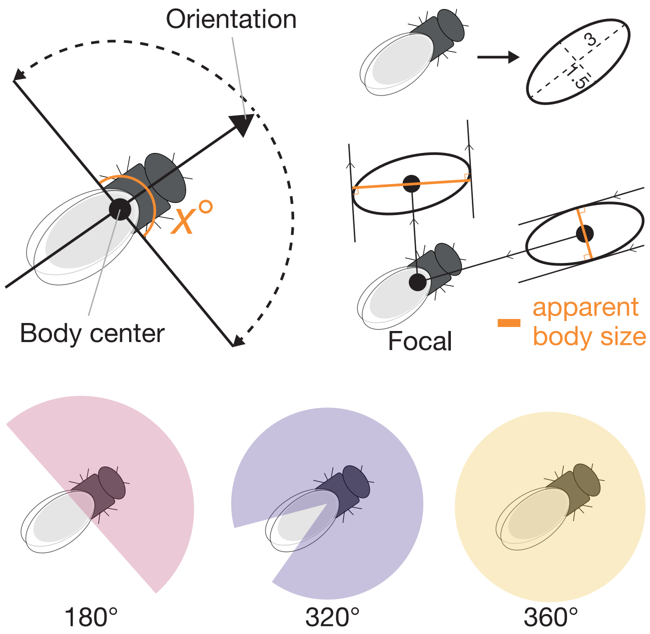


**Figure S5.** The meaning of "*apparent body size*" in visual cue quantification and the method for setting the field of view.

### Dataset in Figshare (separate file).

**df_boot_share.csv**

Mean and within-group variation of movement speed estimated by bootstrapping.

**time_series_share.csv**

Time series of locomotive speed used for the analysis.

**traits_data_share.csv**

Comprehensive trait data for each individual used in the analysis.
